# Identifying the fungal diseases of African Yam Bean (*Sphenostylis stenocarpa* [Hochst ex. A. Rich.] Harms) and their occurrence in South-West Nigeria

**DOI:** 10.1101/2023.09.04.556251

**Authors:** Temitope Oyedele, Iyabo Kehinde, Habeeb Yinka Atanda, Abiodun Oyelakin, Tope Popoola, Luis A. J. Mur

**Affiliations:** Department of Pure and Applied Botany, Federal University of Agriculture, Abeokuta; Department of Life Sciences, Aberystwyth University, United Kingdom

**Keywords:** African Yam Bean, Disease, Fungi, Incidence, Nigeria

## Abstract

*Sphenostylis stenocarpa,* commonly known as African yam bean (AYB), is an orphan crop with high nutritional properties but low yield production due to diseases. Hence, this study accessed the diversity and pathogenicity of fungi associated with AYB in Southwest (SW) Nigeria as a model area of cultivation. The incidence of fungi infecting AYB were surveyed in Oyo, Ondo, Ekiti, Osun, and Ogun states within SW Nigeria during 2018 planting season. The common field symptoms across all sites were tiny spot, brown spot, leaf blight, brown spot with yellow halo, necrotic lesion, and brown spot on pods. A total of 1005 fungi were isolated from leaf and pod samples, and identified morphologically on pure cultures as *Aspergillus* sp*, Botrytis* sp*, Colletotrichum gloeosporioides, Curvularia lunata, Trichoderma harzianum, Macrophomina phaseolina, Pestalotia* sp*, Phoma* sp, *Fusarium verticillioides, F. oxysporum, F. solani*, *Botryodiplodia theobromae*, and *Choanephora curcubitarium and Nigrospora spp. Phoma* sp and *C. gleosporoides* had highest frequency of occurrence 69.9% and 51.9% at early and mature stages, respectively. To conform to Koch’s postulates, the pathogenicities of 12 exemplar strains of the most abundant fungal species were confirmed in controlled glasshouse tests. The identities of *Colletotrichum* sp., *Aspergillus* sp*., Didymella* sp*., Pestalopsis* sp, *Lasiodiplodia theobromae*, *F. solani* and *F. oxysporum* were confirmed based on internal transcribed spacer (ITS) sequencing and comparisons with the Genbank database. AYB germplasm from curated seed banks and farmer donated landraces were grown at the same site in 2020 and identical fungi were isolated. Further, genotypes with reduced disease incidences and incidences were identified. This first study to reveal the diversity of fungi associated with AYB in SW Nigeria that could inform disease management practices.

## 1. Introduction

To meet the human population’s food demand, focus should be on promoting the cultivation and utilization of other crops which have been neglected and underexploited but have the potential to enhance food and nutrition security especially in developing countries. Legumes are reported to have a significant role in improving the nutritional status of a malnourished population particularly in sub-Saharan Africa (Eccles et al., 2019). In the last decade, over three million children have lacked sufficient protein in their diets in Nigerian rural communities. These protein deficiencies can also directly or indirectly affect the health and economic productivity of populations (Ikhajiagbe, 2003). African yam bean (AYB) (*Sphenostylis stenocarpa*) is an underutilized legume and could contribute to countering such protein deficiencies. Further, unlike many legumes AYB is peculiar in that it produces edible seeds as well as underground tubers. The former preferred in West Africa and the latter is used by in East and Central Africa especially among the Bandudus, the Shabas and at Kinshasa in Zaire (Nwokolo, 1987), but there is no intrinsic reason why both seeds and tubers can be used more widely.

AYB is a climbing legume adapted to lowland tropical conditions but can tolerate wide geographical, climatic and edaphic ecologies (Adewale et al., 2008). It thrives on deep, loose sandy and loamy soils with good organic content and good drainage. It grows best in regions where annual rainfall ranges between 800 and 1400 mm, and where temperatures are comprised between 19 and 27 °C (Ecoport, 2009). The wide adaptability of AYB allows it to thrive within the latitudes of 15° North to 15° South and the longitudes of 15° West to 40° East of Africa (Adewale et al., 2008). AYB apparently has its origins in Africa (Potter and Doyle, 1992). The Germplasm Resources Information Network (GRIN, 2009) suggests that areas of diversity include Northeast tropical Africa (i.e., Chad and Ethiopia), East tropical Africa (i.e., Kenya, Tanzania and Uganda), West-Central tropical Africa (i.e., Burundi, Central African Republic and Zaire), West tropical Africa (i.e., Cote d’Ivoire, Ghana, Guinea, Mali, Niger, Nigeria and Togo) and South tropical Africa (i.e. Angola, Malawi, Zambia and Zimbabwe).

The AYB seed coat color varies from white to cream, brown and grey (Kay, 1987, Oshodi et al., 1995). AYB is a good source of protein, carbohydrate, minerals and vitamins. AYB protein content is twice that of potato (*Solanum tuberosum*) and sweet potato (*Ipomoea batatas*) and much higher than those in yam and cassava (Amoatey et al., 2016, Day, 2013). AYB has become an important substitute for the more widely eaten cowpea in areas and a substitute of soya bean in animal feed. The nutritious properties of AYB suggests that it has potential uses in ameliorating malnutrition in many developing countries. This could be by its direct consumption or via fortification and enrichment of less nutritious staples (Ajibola and Olapade, 2016). The presence of amino acids makes AYB useful for fortifying many cereal-based diets that are deficient in protein, and thus addressing the problem of kwashiorkor and marasmus (arising from severe protein malnutrition) among infants. The inclusion of 20% AYB in infant weaning foods can counter the development of such malnutrition associated symptoms in infants (Ijarotimi and Bakere, 2006). Additionally, because of the presence of fiber, some useful starch, and essential fatty acids, AYB is also a good candidate for the development of new functional foods. Medicinally, flavonoids and phenolic acids are two important bioactive compounds that have been identified in AYB (Ade-Omowaye et al., 2015, Oboh and Ademosun, 2006, Oboh et al., 2009, George et al., 2020, Soetan et al., 2018). Indeed, cooked seed can help individuals with diabetic or cardiovascular disorders.

The wider exploitation of AYB is severely limited by plant diseases (Khoury and Makkouk, 2010). Wilting, powdery mildew, and rot gall have been reported to cause severe losses to AYB production (Ameh and Okezie, 2006). However, there is little information on disease occurrence and infective agents that target of AYB. In this work, we survey the symptomology and incidence of disease of AYB in south-western Nigeria and identify some AYB fungal pathogens. Further, we explored the relative resistance to diseases in AYB germplasm obtained from curated seed banks at the International Institute of Tropical Agriculture (IITA, Ibadan, Nigeria) and farmer donated landraces. This information could inform agricultural practice and the development of new disease resistant varieties of AYB.

## 2. Materials and Methods

### 2.1. Field survey, Disease assessment and sample collection

A survey was carried out during 2018 planting season in the AYB growing states of South-west Nigeria, viz: Oyo, Osun, Ondo, Ekiti and Ogun state. In each state, a minimum of two local government sites and two village sites were visited (Fig. 1). The choice of location visited reflected the high production areas of AYB. A total of thirty villages and farmers field were surveyed i.e., one field per village covering all the five of South-western Nigerian. The geographical details of villages surveyed were recorded using Global Positioning Services (GPS) (Table 1).

**Fig. 1.**
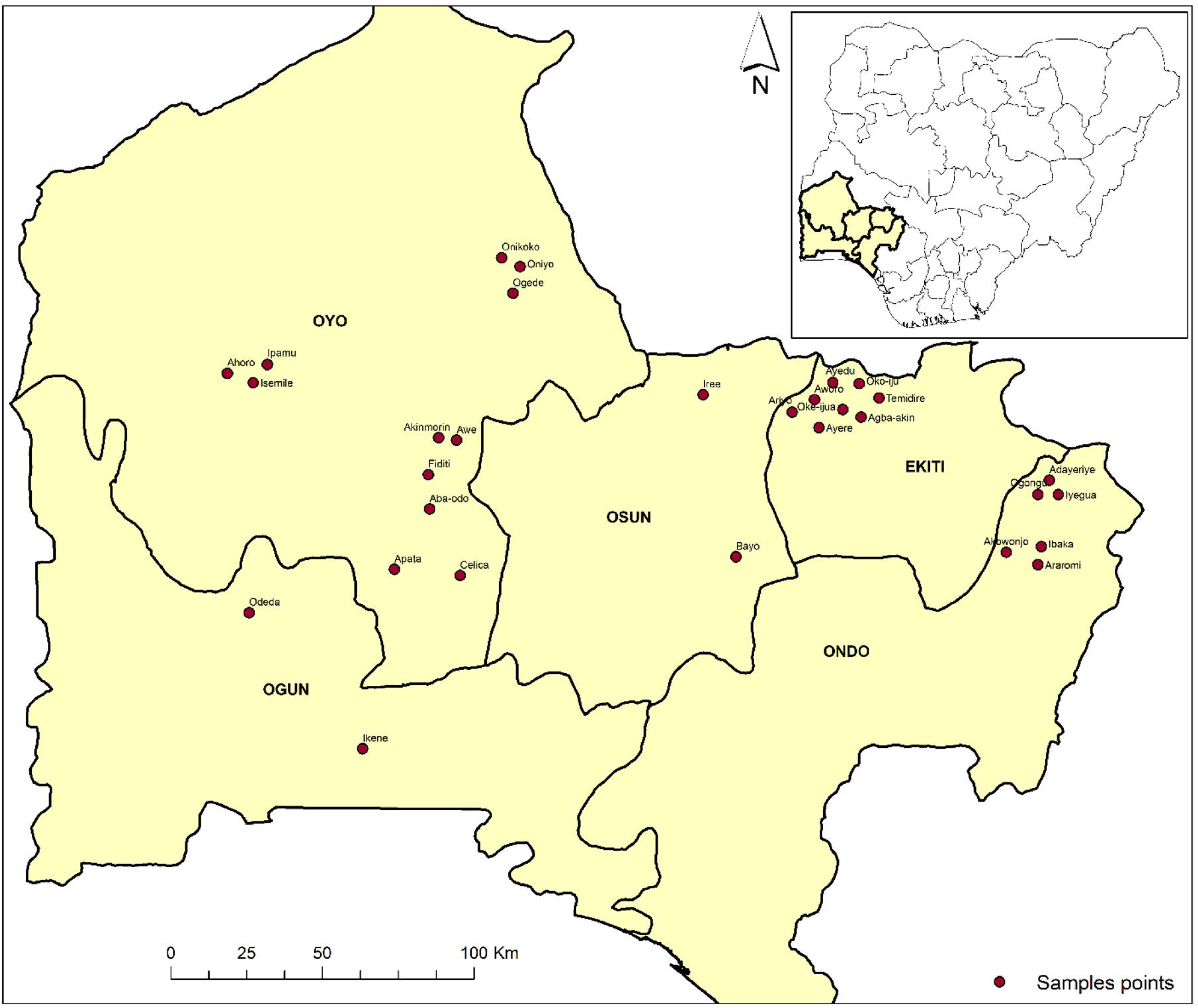
African Yam Bean major growing regions in South-west Nigeria visited in this study

**Table 1.** Sampling location of AYB in South-west Nigeria. KEY: BS; Brown spot, TS; Tiny spot, LB; Leaf blight, BSY; Brown spot with yellow halo, NL; Necrotic lesion, BP; Brown spot.

| Strain | STATE | LG | VILLAGE | LAT | LONG | SYMPTOMS |
| --- | --- | --- | --- | --- | --- | --- |
| 1 | OYO | Akinyele | Aba-odo | 7.575945 | 3.912677 | LB,BS,BSY,NL, BP, |
| 2 |  | Egbeda | Celica | 7.37898763 | 4.0031447 | BS, LB, BP, BSY |
| 3 |  | Afijio | Fiditi | 7.67796752 | 3.90860196 | BS,TS,BP,NL |
| 4 |  | Ido | Apata | 7.37339097 | 3.84009497 | BS, TS, LB,NL |
| 5 |  | Kajola | Ipamu | 7.99651146 | 3.46336712 | BSY,TS,NL |
| 6 |  | Afijio | Awe | 7.78031767 | 3.99291704 | TS,BSY,NL,BP |
| 7 |  | kajola | Ahoro | 7.9943927 | 3.39167367 | BS,TS,NL |
| 8 |  | Kajola | Akinmori | 7.78749874 | 3.93943267 | BS, LB,NL,BS |
| 9 |  | kajola | Isemile | 7.99414472 | 3.3892045 | BS, TS, NL |
| 10 |  | Orire | Ogede | 8.28606977 | 4.1870379 | TS, BS, NL |
| 11 |  | orire | Onikoko | 8.30173355 | 4.16568791 | BS, LB,NL |
| 12 |  | orire | Oniyo | 8.29507768 | 4.18053492 | BS, BSY, BS |
| 13 | EKITI | Ijero | Aworo | 7.92238585 | 5.11665955 | TS, LB, NL |
| 14 |  | Ido-osi | Ayedu | 7.9197948 | 5.1338453 | TS, BS, NL |
| 15 |  | Ijero | Ayere | 7.89581028 | 5.1295921 | LB, BS, NL |
| 16 |  | Ido-osi | Agba-akin | 7.91451123 | 5.16794369 | BS,TS, BP |
| 17 |  | Ido-osi | Oke-ijua | 7.91217035 | 5.16349416 | TS, BS,BP,NL |
| 18 |  | Ijero | Ariyo | 7.92717835 | 5.10094983 | BS,TS, BP |
| 19 |  | Ilejemeje | Oko-iju | 7.92756381 | 5.17334244 | TS, BS, BSY, BP |
| 20 |  | Ilejemeje | Temidire | 7.92467575 | 5.18273521 | LB,TS, BP |
| 21 | ONDO | Akok SW | Akowonjo | 7.44797954 | 5.62276808 | BS, LB,NL |
| 22 |  | Akok SW | Araromi | 7.45670259 | 5.72188083 | TS,BS, BP |
| 23 |  | AkokNW | Adayeriye | 7.64616211 | 5.74210298 | BS,BSY,.BP |
| 24 |  | AkokNW | Iyegua | 7.63858899 | 5.74369429 | TS, BS, BP |
| 25 |  | AkokNW | Ogongu | 7.63967634 | 5.74595921 | BS,BP |
| 26 |  | Akok SW | Ibaka | 7.464472 | 5.72626343 | BS,TS,NL |
| 27 | OSUN | Atakom E | Bayo | 7.43419345 | 4.82114978 | LB,TS, NL |
| 28 |  | Boripe | Iree | 7.92684229 | 4.7161169 | TS, BS,NL,BP |
| 29 | OGUN | Ikene | Ikene | 6.865 | 3.7143 | BS,BP |
| 30 |  | FUUNAB | Odeda | 7.2437 | 3.3433 | BS,TS, BP |

At the first visit 20 plants were tagged per field at the middle and edge, and assessment of infected plant tissues commenced 6 weeks after planting. This represented the “early” assessment of disease incidence. The plots were revisited at “maturity” (at 20 weeks) when disease incidence on the leaves and pods were assessed. Then, each diseased leaf, and pods samples were collected to isolate and identify the casual organisms. Diseases plants were imaged, and the infected parts were promptly collected, into labeled polythene bags and taken to the pathology laboratory.

Different types of symptoms observed on diseased AYB from the field survey were recorded and the disease incidence were determined.

The proportion of diseased plants were estimated by:

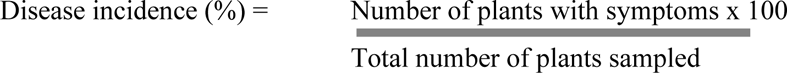

Follow-up assessments of disease incidence were conducted in early June, 2020 at the experimental fields in the Department of Pure and Applied Botany at the Federal University of Agriculture, Abeokuta (Latitude 7.2437, longitude 3.3433, elevation of 165m above sea level) with a mean annual rainfall of 1400 mm. The environment is considered as derived savanna with mean annual temperature of between 22.7°C −34.5 °C. The germplasm used (n=40) included AYB lines obtained from IITA and landraces grown typically across the previously sampled areas (Fig. 1). The experiment was laid out in Randomized Complete Block Design (RCBD) in three replicates. Three seeds were planted on 5m long ridges of loamy soil in single row plots with spacing of 1.0m x 0.5m between rows and within rows. The plants were later thinned to one plant per ridge, watered regularly and weeds were regularly rouged off.

### 2.2. Isolation of fungi from samples collected

The infected tissues were cut alongside healthy tissues with a sterile scalpel. In the laboratory, diseased leaf and pod parts were surface sterilized in 1% sodium hypochlorite (NaOCl) for 10 min and rinsed in three changes of sterile distilled water. Five segments of diseased parts were placed on each Petri dish on Potato Dextrose Agar (PDA) culture medium. The plates were incubated for 5 to 7 days at room temperature 28 ±2°C and observed daily from the 2^nd^ day. Fungal colonies were subcultured on PDA until pure cultures are obtained. Cultural characteristics such as colony appearances, mycelial textures, and pigmentations on both obverse and reverse on PDA plates were observed after 5-7 days of incubation.

### 2.3. Percentage frequency of occurrence

Occurrence of fungi was determined by counting the number of times each individual fungus occurred divided by the total number of fungi and expressed as a percentage. Thus.

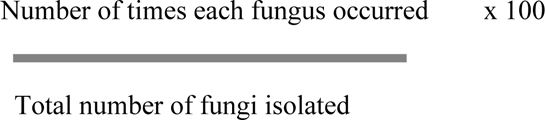

### 2.4. Fungal identification based on morphology

Macroscopic characteristics involved examination of fungal colonies, pigmentation, and conidial masses from 7-day-old PDA cultures. Identification was performed by observing the culture plates and microscopic structures under a light microscope.

### 2.5. DNA extraction, PCR amplification, sequencing and identification of species

Genomic DNA from the isolated fungi was conducted using CTAB Protocol described by (Carter-House et al., 2020). Primers ITS1 (5′-TCCGTAGGTGAACCTGCG-3′), and ITS4 (5′-TCCTCCGCTTATTGATATGC-3′) were used to amplify ribosomal internal transcribed spacer (ITS). The polymerase chain reaction was carried out at 94 °C for 5 min, followed by a 35 cycle 3 step PCR, each cycle included a denaturing step at 94 °C for 30 sec, an annealing step at 55 °C for 30 sec, and an extension step at 72 °C for 30 sec, and finally kept at 72 °C for 10 min for final extension. Extracted and amplified DNA fragments were visualized on a 2 % agarose gel in Tris-acetate-EDTA buffer, stained with ethidium bromide, at 60 V for 50 min and visualized under a UV trans-illuminator.

The PCR products were Sanger sequenced at the Translation Genomic laboratory at the Institute of Biological, Environmental and Rural Sciences, Aberystwyth University. The sequences generated in this study was compiled and edited using Bio-Edit sequence alignment editor (version 7.2.5) and compared with sequences from the GenBank database using the National Center for Biotechnology Information (NCBI) Basic Local Alignment Search Tool (BLAST). Multiple alignments of nucleic acid and amino acid sequences were performed using CLUSTALW (Thompson et al., 1994). The Genbank accession numbers for the fungal ITS sequences are listed in Table 2.

**Table 2.** Fungal strains isolated from African Yam Bean identified by internal transcribed spacer (ITS) sequencing.

| Isolate | Accession number | Strain ID |
| --- | --- | --- |
| 1 | OR362334 | *Fusarium oxysporum* |
| 2 | OR362335 | *Fusarium solani* |
| 3 | OR362336 | *Fusarium oxysporum* |
| 4 | OR362337 | *Aspergillus* sp. |
| 5 | OR362338 | *Fusarium solani* |
| 6 | OR362339 | *Fusarium solani* |
| 8 | OR362340 | *Lasiodiplodia* sp. |
| 11 | OR362342 | *Didymella* sp. |
| 14 | OR362341 | *Pestalotiopsis* sp |
| 15 | OR362343 | *Colletotrichum* sp. |

The phylogenetic relationships of the amplified ITS region sequences were constructed by the Maximum likelihood method as implemented in MEGA v10.1.8 (Kumar et al., 2018) with 1000 random bootstrap replications. *Meyerozyma guilliermondii* strain APMSU5 was used as an out-group.

### 2.6. Controlled environment pathogenicity testing of the isolated organisms on healthy AYB

A total of twelve fungal isolates obtained in this study were tested for pathogenicity on healthy leaves. Seeds were sown in a pot containing three and half (3.5 kg) sterile sandy loamy soil placed in greenhouse of the International Institute of Tropical Agriculture (IITA) Ibadan (Nigeria). All the fungi isolated from different symptoms AYB leaf and pod were inoculated into healthy AYB plant to determine whether they could induce similar symptoms on re-inoculation. Fungal suspension was prepared from the 10 days old culture plates of the isolated fungi. The mycelia mass of the fungus growth culture in the Petri dishes were scooped out into a sterile conical flask, which contains 10 mL of sterile distilled water, and a drop of Tween 20 detergent (for spore dispersal) was added (Kehinde, 2008). The resulted suspension of each isolate was adjusted to 10^5^ spore /ml. Thereafter, seedlings were inoculated by spraying on leaves by using hand sprayer (Han *et al*., 2003) and were covered with transparent polythene bag for 24 h to maintain high humidity. The disease symptoms were observed up to 70 days. Mixture of 4 isolates were also made to made up a cocktail (CKL) of strains to ease plant screening. Control plants were sprayed with sterile distilled water for comparison. Disease assessment was a visual observation of presence or absence of symptoms after inoculation. Fungi were reisolated from the diseased leaf and morphological compared with original isolates.

### 2.7. Severity index

The severity index of the disease described the damage caused by the diseases on plants leaves and pod. Method of Nakawuka and Adipala 1997 was used to score the diseased plants as follows: 1= 0% infection (no visible symptoms on either leaves or pods), 2 = less than 10% of infection (scattered lesions on either leaves or pods), 3 = 10%–20% infection (extensive spotting of leaves or pods), 4 = 20%–50% infection (lesions coalescing covering half of leaves or pods), 5 = more than 50% infection (leaves or pods severely damaged).

## 3. RESULTS

### 3.1 Surveying AYB across SW Nigeria

Thirty villages in AYB growing states of Southwest (SW) Nigeria were surveyed for diseases of AYB (Fig. 1). The observed symptoms ranged from, tiny spot symptoms appeared as small dark spots with grey centre on leaf blades (Fig. 2a); brown spot appeared as a scattered brown irregular spots that coalesced together (Fig. 2b); brown spots with a yellow halo surrounding each spot, the centre of the spot being dry to give a grey colour (Fig. 2c); leaf blight gives a brown lesion on leaf tips and vein (Fig. 2d); necrotic lesion on the pod with brown or black color, which eventually enlarged to cover almost the whole pod (Fig. 2e) and brown spot on pod appears as specks of pycnidia showing fruiting bodies on senescing pods with seeds showing cracked and shriveled seed coats (Fig. 2f).

**Fig. 2.**
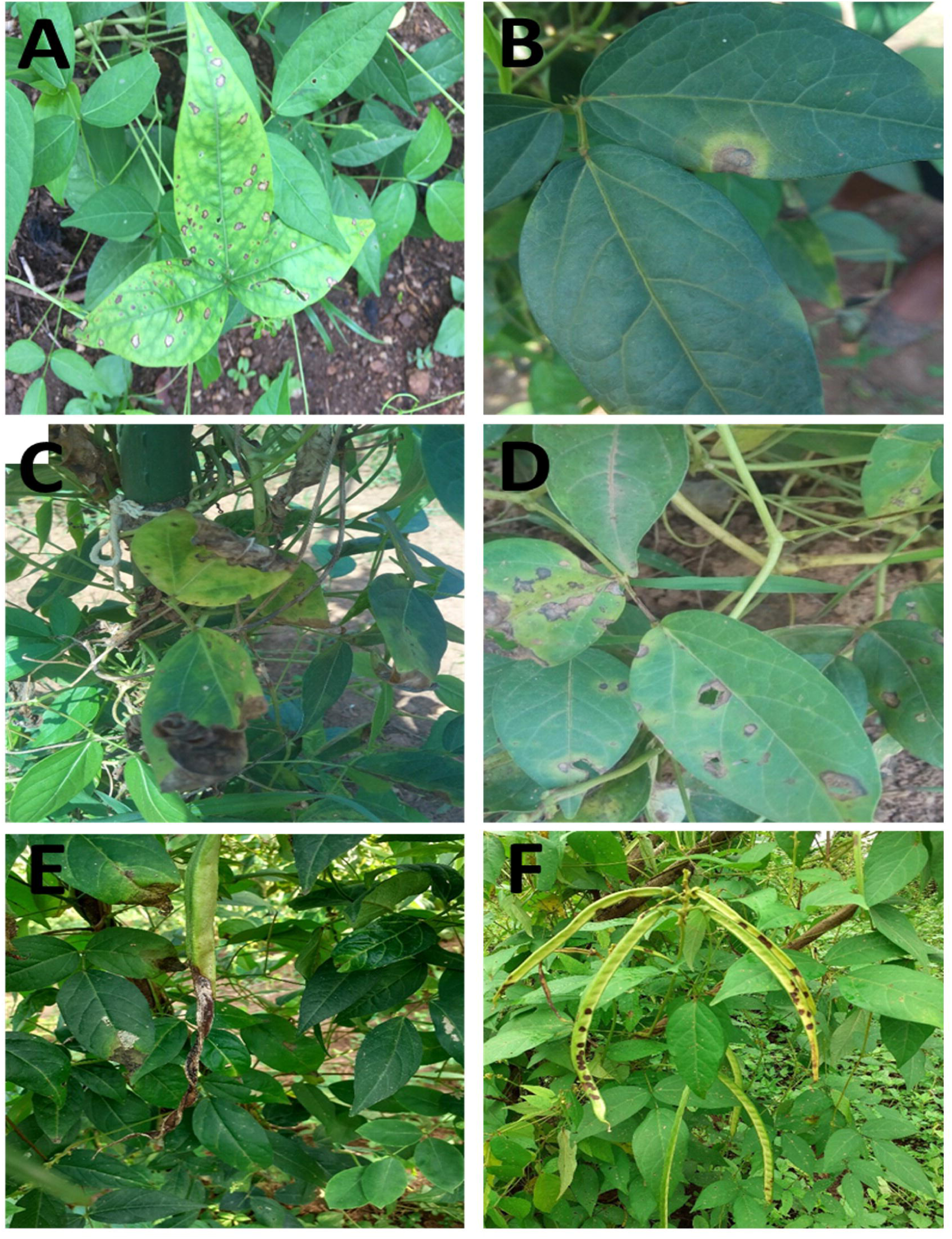
Different symptoms produced by Fungi in AYB leaf and pod under field conditions **(A)** tiny spot symptoms with small dark spots with grey centre on leaf blades, **(B)** brown spot scattered brown irregular spots that coalesced together, **(C)** brown spots with a yellow halo surrounding each spot, the centre of the spot being dry to give a grey colour, **(D)** leaf blight gives a brown lesion on leaf tips and vein, **(E)** necrotic lesion on the pod with brown or black colour, which eventually enlarged to cover almost the whole pod and, **(F)** brown spot on pod appears as specks of pycnidia showing fruiting bodies on senescing pods with seeds showing cracked and shrivelled seed coats.

Disease incidence at the sites was recorded at two visits; at 6 weeks post AYB sowing (6 weeks “early”) and at maturity (20 weeks) when pods were formed (Fig. 3). At the early stage, incidence varied from 25 - 55 % in different fields of Oyo state. The lowest diseases incidence (25 %) was observed in Onikoko. The highest (55 %) was observed in Akinmorin, Fiditi and Abaodo in Oyo state. AYB plants with symptoms varied from 5 - 45% in different fields of Ekiti state with the lowest (5 %) observed in Ayere. In Ondo, the diseases incidence ranges from 10 - 50% with the highest incidence in (50 %). Disease incidence varied from 30- 40% in Ogun state. The highest incidence was observed in Osun state at > 50 %. Diseases incidences were greater at mature rather than early stage in all the villages visited. The highest diseases incidence recorded at maturity stage on leaf was in Fiditi (84 %) while the least was observed in Agba-akin and Iyegua (20 %). The highest diseases incidence on pod was recorded in Celica (55 %) in Oyo state.

**Fig. 3.**
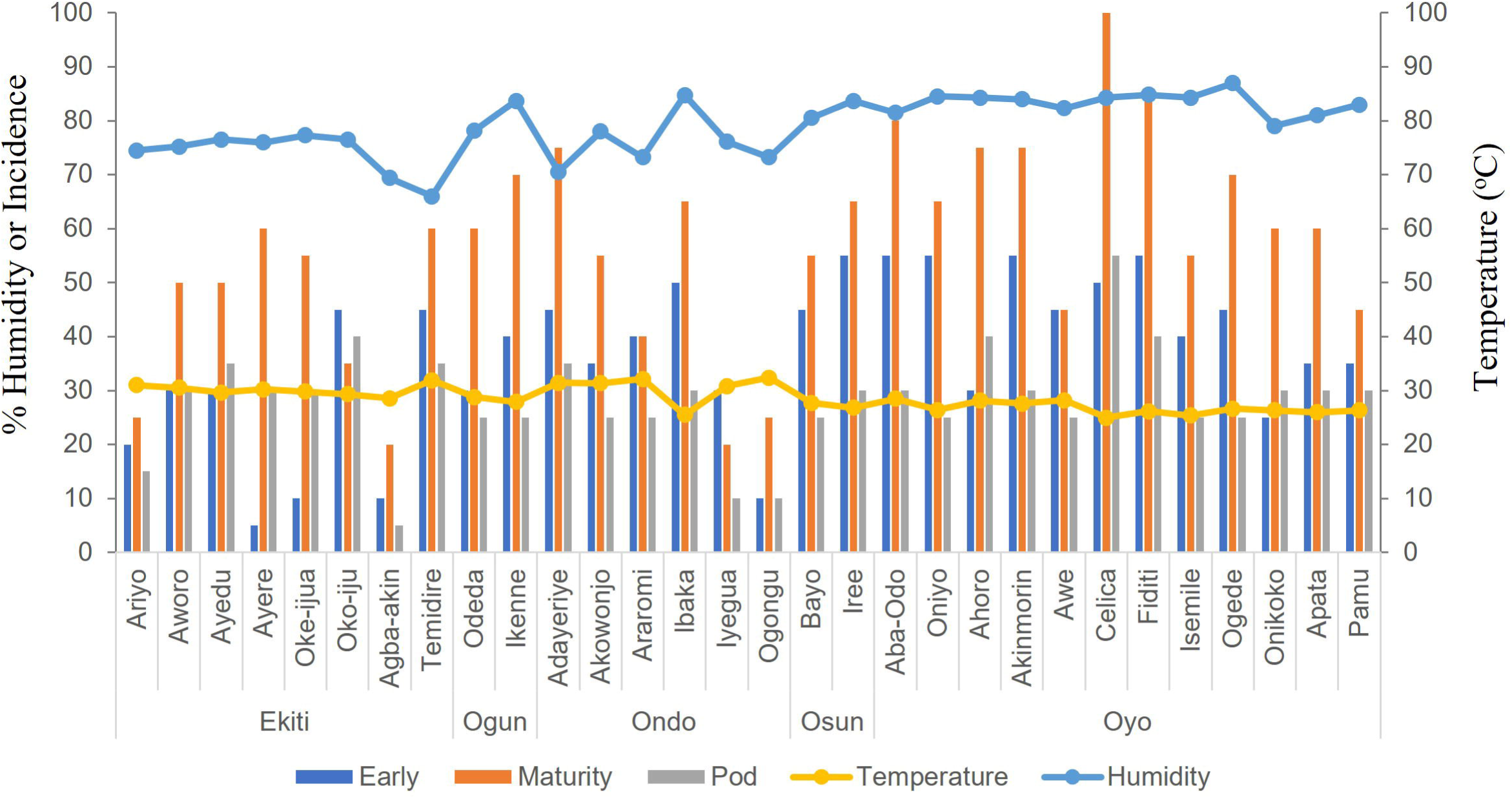
Percentage disease incidence of AYB in the states Southwest Nigeria related to humidity and temperature

At the mature stage, infected leaves and pods were sampled. The associated fungal strains were isolated to homogeneity by successive rounds of sub-culture. A total of 1005 isolates of fungi were isolated, and cultures were identified based on morphological characters and categorized into genera. The occurrence of identified fungi associated with each symptom and growing stages (early and maturity) of AYB is shown in Fig. S1. These observational data are graphically displayed in as a circos plot (Fig. 4) where particularly *Phoma* spp (82.2 %) but also *Fusarium verticolloides* (13.1%) *Cuvularia lunata* (1.0%) were predominant in “tiny spot” symptoms. With necrotic lesions on pod *Colletotrichum gleosporoides* showed a high prevalence (64.1 %), however others such as *Phoma* spp (8.0%), *Macrophomina phaseolina* (4.0%)*, Fusarium oxysporum* (0.6%), and *Fusarium solani (*0.6%) (Fig. 4). Furthermore, *Phoma* spp had the highest percentage occurrence (64.6%) (81.3%) on brown spot-on leaf and pod respectively. When the sites of infection on AYB plants were considered (Fig. S2), A*spergillus* sp*, Botrytis sp, Colletotrichum gloesporoides, Macrophomina phaseolina, Pestalotia sp, Phoma* sp *Fusarium verticolloides, Fusarium oxysporum* and *Botrydipolodia theobromae* were isolated at early (leafy) stage of AYB development. The most significant occurrences at both early and maturity stages were *Phoma* spp (69.9 %) and *C. gleosporoides* (51.9 %), respectively. Considering the geographical distribution of the isolated strains (Fig. 5) *Phoma* spp and *C. gleosporoides* were particularly prevalent in Oyo, Osun, Ogun, Ekiti and Ondo States. There were low occurrences of *Pestalotia* sp in Ekiti, Ondo and Oyo states. It should be noted that every isolated pathogen was identified in every state.

**Fig. 4.**
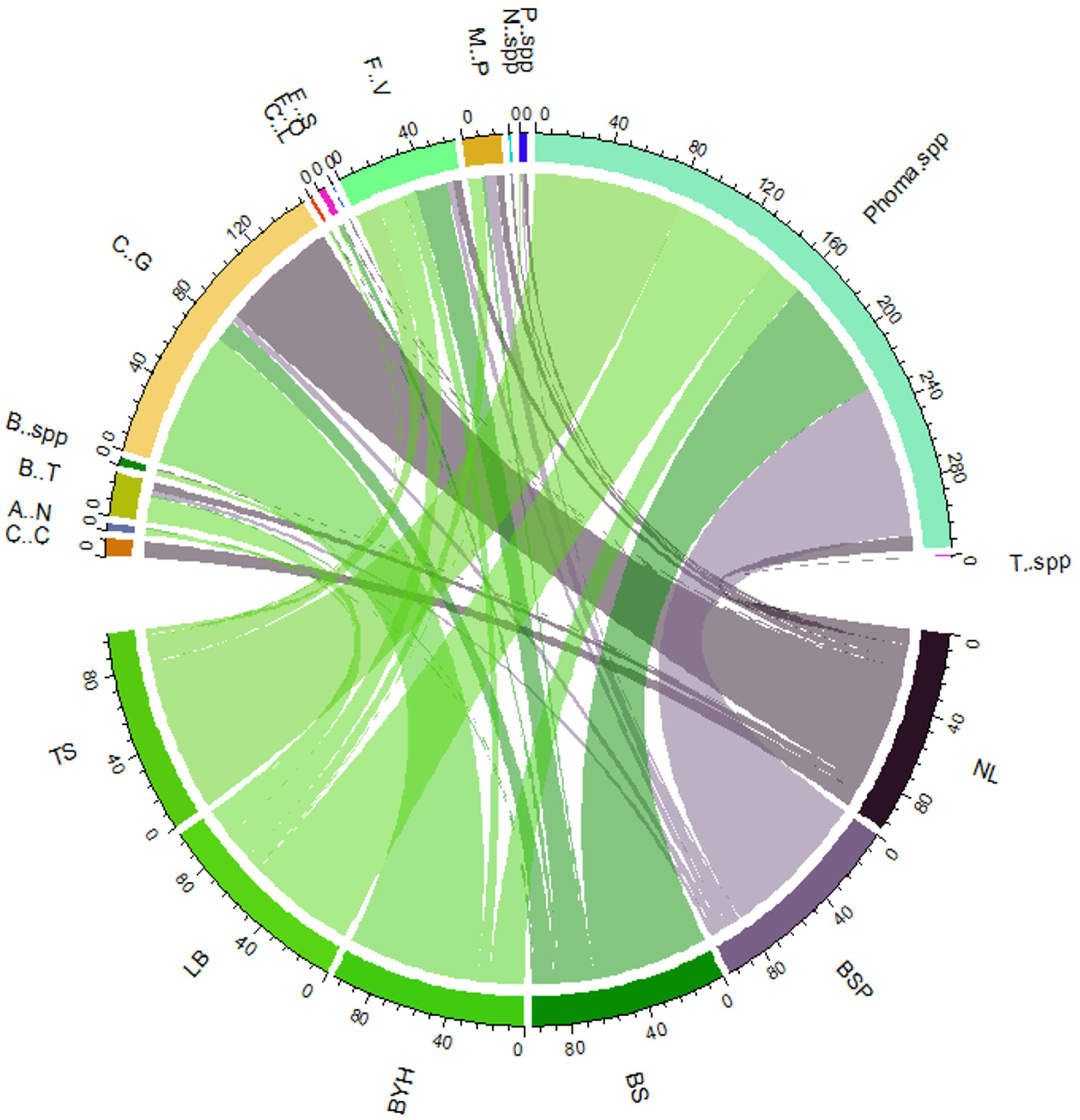
Fungi species linked to symptoms on African Yam Bean Symptom key: TS - Tiny spot; LB- Leaf blight; BYH- Brown spot with yellow halo; BS- Brown spot; BSP- Brown spot on pod; NL- Brown lesion on pod. Pathogen Key: C.C- *Choanephora cucurbitarium*, T.SPP- Trichoderma spp, C.L- *Curvularia lunat*a, A.N-A*spergillus* sp*, B.SPP-Botrytis sp,C.G- Collectrotricum gloesporoides, M.P-Macrophomina phaseolina, P.SPP-Pestalotia sp, Phoma* sp, F.V- *Fusarium verticolloides, F.O-F. oxysporum, F.S- F. solani, N.SPP-Nigrospora spp* and B.T- *Botrydipolodia theobromae*

**Fig. 5.**
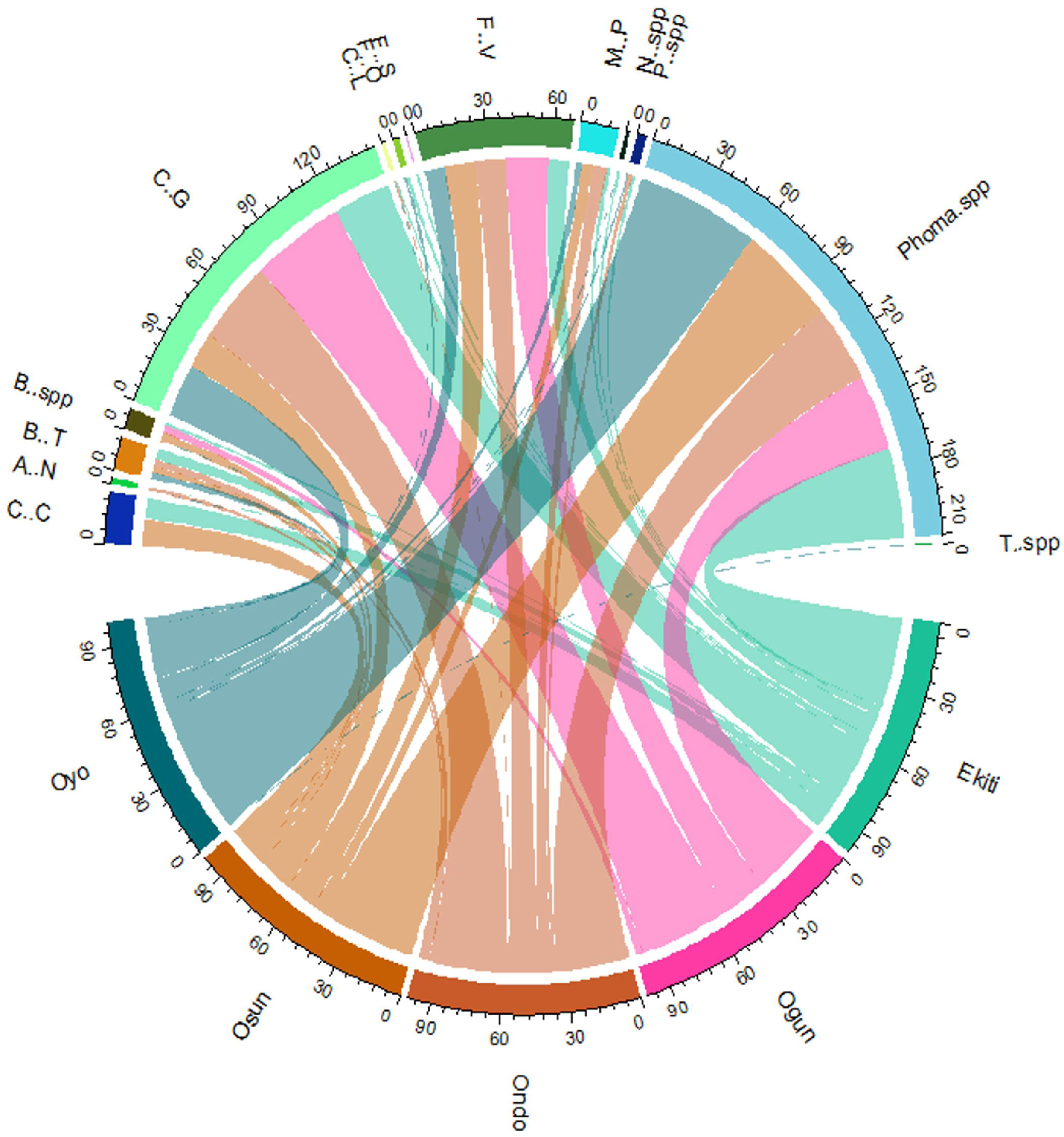
The occurrence (%) of fungal species isolated from African Yam Bean in Southwest Nigerian States Pathogen Key: C.C- *Choanephora cucurbitarium*, T.SPP- Trichoderma spp, C.L- *Curvularia lunat*a, A.N-A*spergillus* sp*, B.SPP-Botrytis sp, C.G- Collectrotricum gloesporoides, M.P-Macrophomina phaseolina, P.SPP-Pestalotia sp, Phoma* sp, F.V- *Fusarium verticolloides, F.O -F. oxysporum, F.S- F. solani, N.SPP-Nigrospora spp* and B.T- *Botrydipolodia theobromae*

### 3.2. A field assessment of disease incidence and severity on AYB germplasm

The survey of field sites (Fig. 1) allowed us to establish a collection of farmers’ AYB landraces that were commonly grown in that region. This led to our collecting 20 landraces designated “FSS”. Improved AYB cultivars lodged at IITA were also obtained and were designated “TSS” lines. As each pathogen identified was found in each SW Nigerian region (Fig. 5) a single field sites was used to assess the responses of the difference lines to disease. At 20 weeks following sowing all genotypes showed disease that matched the previously observed symptoms. Fungal strains were isolated from this experiment which were identified based on morphologies. The identified fungal strains aligned with those observed in the previous survey over the SW Nigerian region (Table S1). Therefore, the field experiment in 2020 can be considered to validate the previous 2018 survey. However, different AYB genotypes exhibited but at varying incidences (Fig. S2A) and severities (Fig. S2B). These data were used to derive a dendrogram grouping genotypes according to their field responses to disease (Fig. 6).

**Fig. 6.**
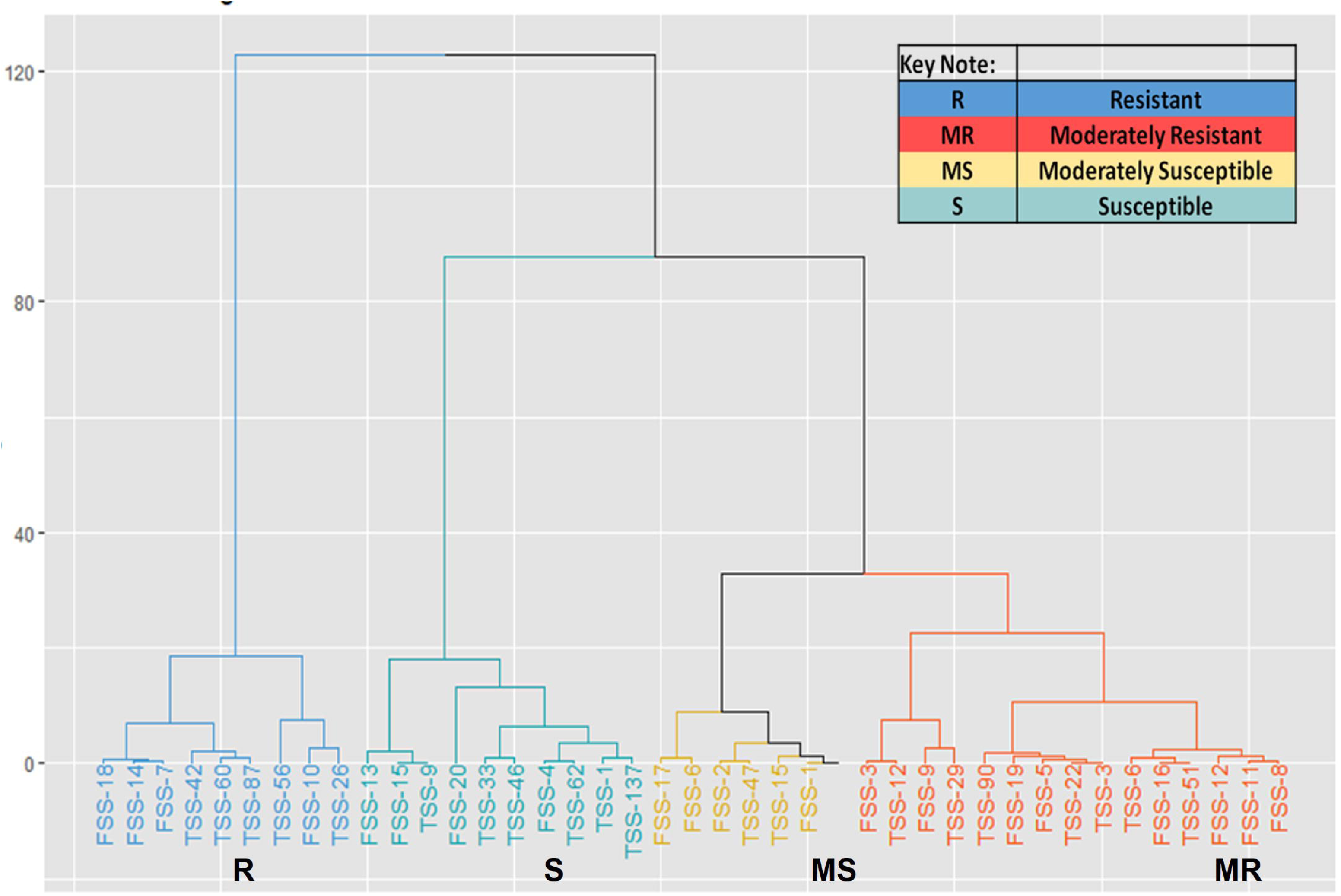
Dendrogram grouping African Yam Bean farmers landrace (FSS) and cultivar (TSS) based on their responses to disease in the field.

Nine of the AYB accessions were resistant to the diseases and these included four farmers’ landraces (FSS 18, 14, 7 and 10) and five IITA accessions (TSS 42, 60, 87, 56 and 26). 10 genotypes including four farmers’ landraces (FSS 13, 15, 20 and 4) and six IITA accessions (TSS 9, 1, 33, 137, 46 and 62) were susceptible to the disease. Six genotypes comprising of four landraces (FSS 16, 6, 2 and 1) and two IITA lines (TSS 47 and 16) were moderately susceptible to the disease. The remaining fifteen genotypes comprising of landraces (FSS 3, 9, 19, 5, 16, 12, 11 and 8) and seven IITA lines (TSS 12, 29, 90, 22, 3, 6 and 51) were moderately resistant.

### 3.3. Genetic confirmation of the identities of some fungi isolated from AYB

Certain fungal strains were distinctive in their culture morphologies, color and mycelial growth patterns. Of these DNA was isolated from seven isolates and the ITS region (ITS1-ITS4) was amplified to yield fragments ranging between 550 to the 600 bp. The amplicons were pair-end sequenced and the consensus sequence assessed by BLAST analysis. Amplicon sequences and the ITS sequences of matching voucher species were used to derive phylogenetic trees using the maximum likelihood method. The tree was out grouped by *Meyerozyma guilliermondii* (Fig. 7). As only a single locus was sequenced, we were confident of identifications down to genus-levels. In most cases, the ITS sequencing confirmed the morphological based identification of the strains derived from AYB. In case of the presumptive *Phoma* spp. isolate, this was indicated to be a sister genera, *Didymella.* within the *Didymellaceae*.

**Fig. 7.**
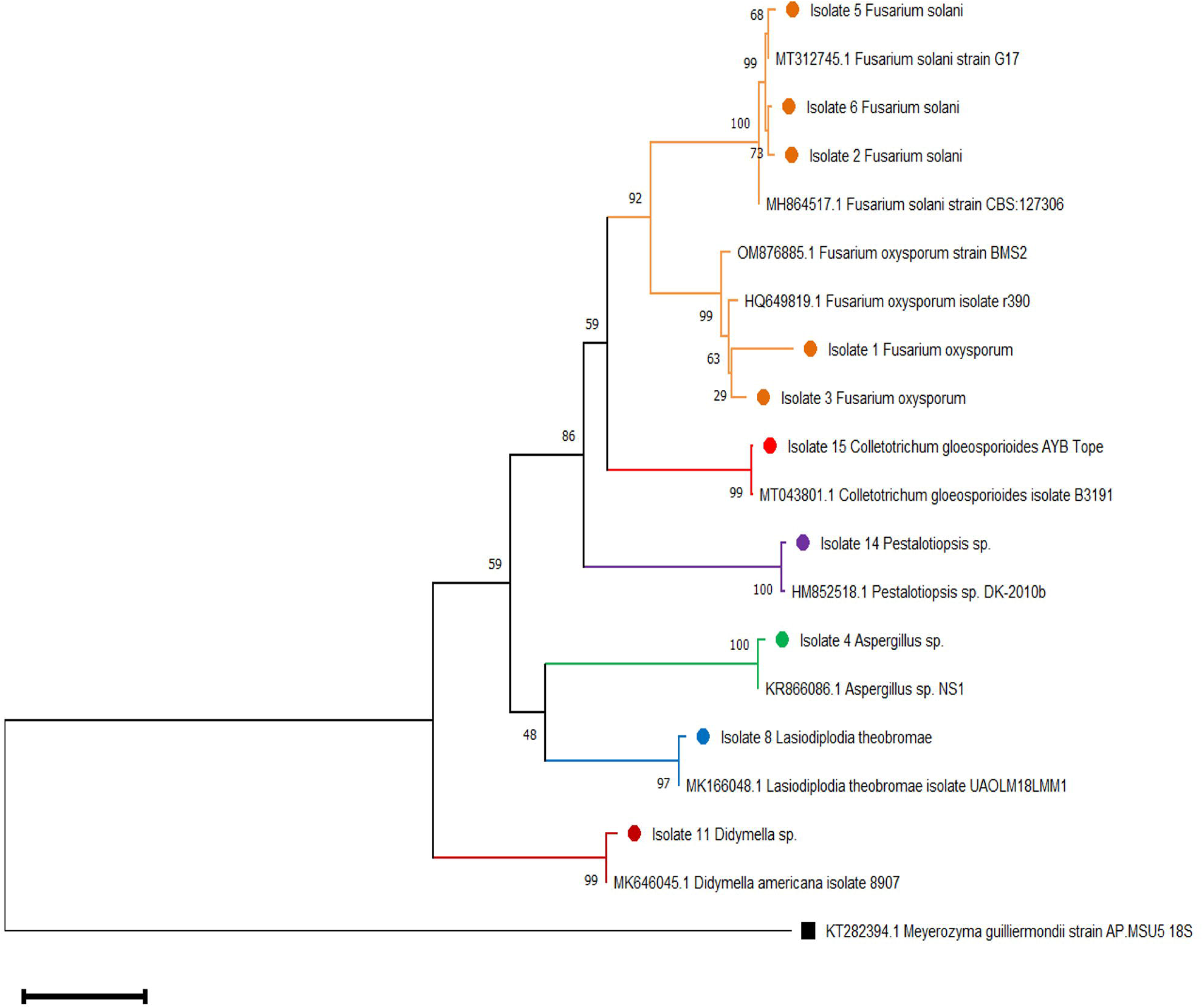
Phylogenetic analysis of the fungal species isolated from African Yam Bean based on internal spacer (ITS) sequence variation.

### 3.4. Pathogenicity testing

In accordance with Koch’s postulates, we sought to confirm that the isolated strains were pathogenic by reinfection. A mixture of *Didymella sp, Colletotrichum gloeosporioides, Fusarium oxysporum and Botryodiplodia theobromae* made up the cocktail (CKL) was also used as a means to rapidly screen the fungal strains and also assess synergistic effects in infection. Inoculation of AYB seedlings with the identified fungi resulted in the production of brown spots of various sizes, blight and spots with yellow halo on leaf. Analysis of variance showed that all the parameters evaluated have significant effect on the disease severity (*P*<0.001), interaction between the accession and isolate; accession and days and isolate x days were also significant (*P* ≤ 0.001.) (Table 3). All the fungal isolates tested were aggressive on AYB showing high severity scores that are significantly different (*P*<0.05) from the control. Four isolates showed 100% incidence rates on plants of AYB accession TSS 137. *Trichoderma* was weakly aggressive on all the accession showing incidence ranging from 0 – 7.4 % (Table 4). *Didymella* had the highest disease severity across all of the tested AYB accessions (3.2 – 4.1) whilst, unsurprisingly, as an endosymbiont, *Trichoderma* had the least mean severity score (1.0 – 1.1) (Table 5). Most of the isolates showed average mean disease severity above 3.0. The percentage incidence and the mean severity score of the disease increase continuously for each day (DAI) across all the accession under this study with the 70 DAI showing the highest severity scores (4.5) and percentage incidence (92.5%) (Table 5). AYB accession TSS 62 exhibited the least mean severity score, whilst TSS 137 showed highest mean severity. CKL infections exhibited no evidence of synergistic effects on the disease parameters.

**Table 3.** Diseases severity scores on AYB lines.

|  | Df | Sum<br>square | Mean<br>square | F value | Pr(>F) |
| --- | --- | --- | --- | --- | --- |
| Accession | 5 | 34.96 | 6.99 | 34.78 | $< 2.2 \times 10^{-16}$ *** |
| Isolate | 13 | 1713.51 | 131.81 | 655.58 | $< 2.2 \times 10^{-16}$ *** |
| DAI | 8 | 2306.83 | 288.35 | 1434.18 | $< 2.2 \times 10^{-16}$ *** |
| Accession &<br>Isolate | 65 | 117.63 | 1.81 | 9 | $< 2.2 \times 10^{-16}$ *** |
| Accession & DAI | 40 | 35.46 | 0.89 | 4.41 | $< 2.2 \times 10^{-16}$ *** |
| Isolate & DAI | 104 | 372.81 | 3.59 | 17.83 | $< 2.2 \times 10^{-16}$ *** |
| Accession &<br>Isolate & DAI | 520 | 130.46 | 0.25 | 1.25 | 0.0008337 *** |
| Residuals | 1512 | 304 | 0.2 |  |  |

Significance codes: '\*\*\*' $P = 0.001$ , '\*\*' $P = 0.01$ , '\*' $P = 0.05$ "

**Table 4.**
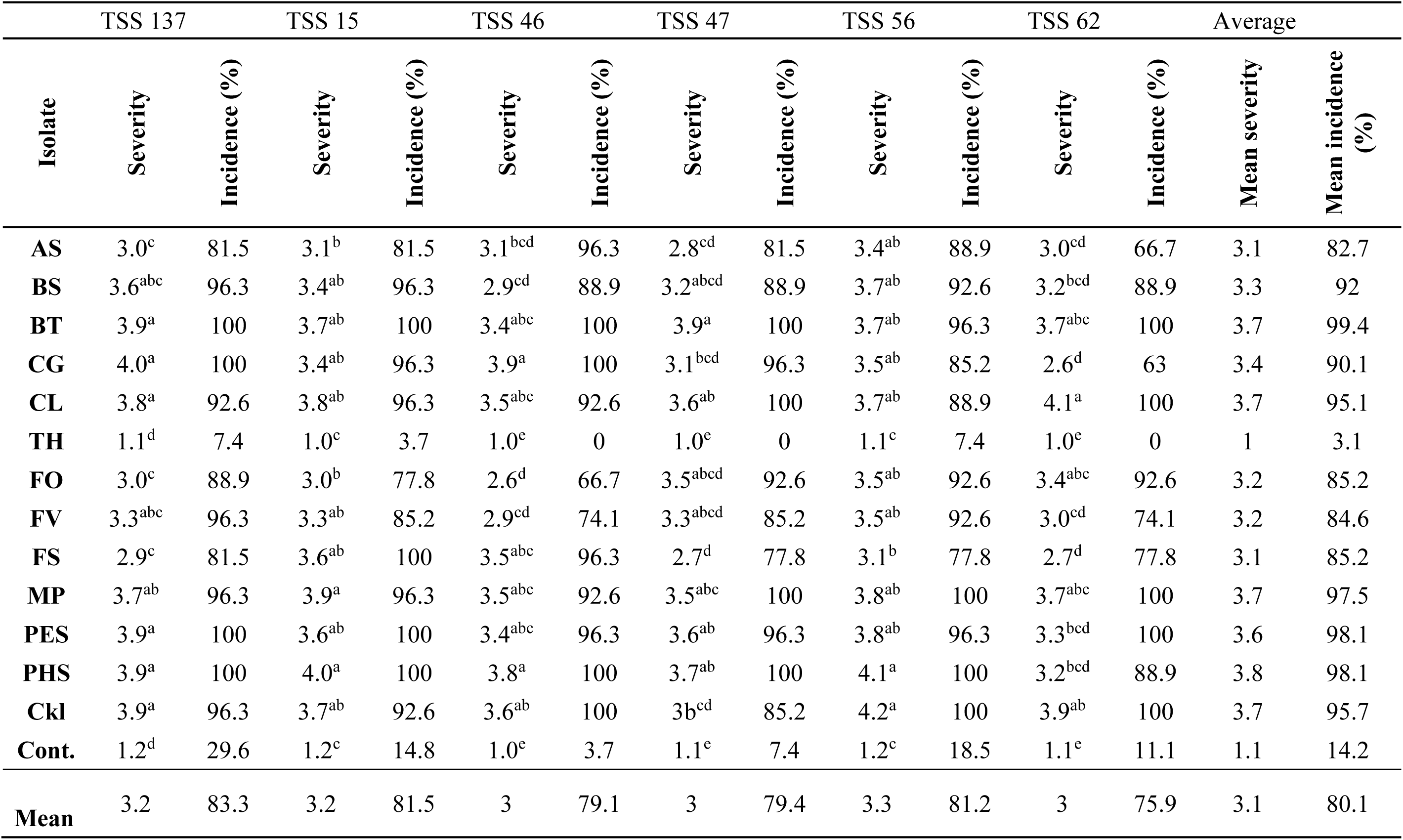
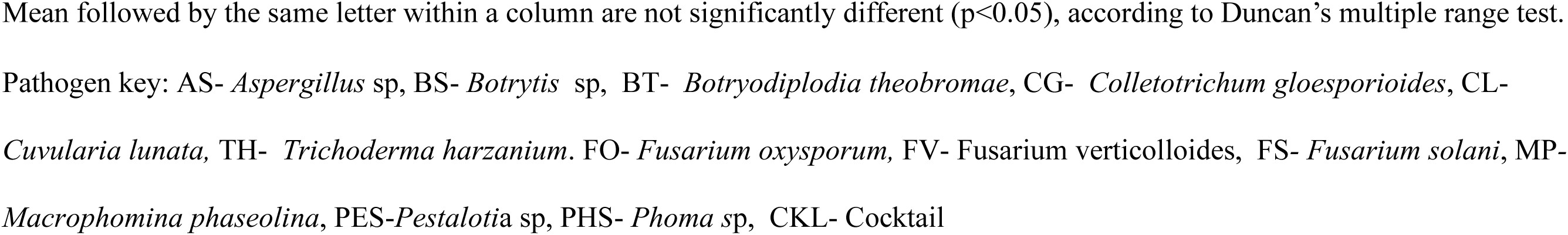
Disease severity and percentage incidence on African yam bean inoculated with fungi isolates under greenhouse conditions.

|  | TSS 137 |  | TSS 15 |  | TSS 46 |  | TSS 47 |  | TSS 56 |  | TSS 62 |  | Average |  |
| --- | --- | --- | --- | --- | --- | --- | --- | --- | --- | --- | --- | --- | --- | --- |
| **Isolate** | **Severity** | **Incidence (%)** | **Severity** | **Incidence (%)** | **Severity** | **Incidence (%)** | **Severity** | **Incidence (%)** | **Severity** | **Incidence (%)** | **Severity** | **Incidence (%)** | **Mean severity** | **Mean incidence (%)** |
| **AS** | 3.0<sup>c</sup> | 81.5 | 3.1<sup>b</sup> | 81.5 | 3.1<sup>bcd</sup> | 96.3 | 2.8<sup>cd</sup> | 81.5 | 3.4<sup>ab</sup> | 88.9 | 3.0<sup>cd</sup> | 66.7 | 3.1 | 82.7 |
| **BS** | 3.6<sup>abc</sup> | 96.3 | 3.4<sup>ab</sup> | 96.3 | 2.9<sup>cd</sup> | 88.9 | 3.2<sup>abcd</sup> | 88.9 | 3.7<sup>ab</sup> | 92.6 | 3.2<sup>bcd</sup> | 88.9 | 3.3 | 92 |
| **BT** | 3.9<sup>a</sup> | 100 | 3.7<sup>ab</sup> | 100 | 3.4<sup>abc</sup> | 100 | 3.9<sup>a</sup> | 100 | 3.7<sup>ab</sup> | 96.3 | 3.7<sup>abc</sup> | 100 | 3.7 | 99.4 |
| **CG** | 4.0<sup>a</sup> | 100 | 3.4<sup>ab</sup> | 96.3 | 3.9<sup>a</sup> | 100 | 3.1<sup>bcd</sup> | 96.3 | 3.5<sup>ab</sup> | 85.2 | 2.6<sup>d</sup> | 63 | 3.4 | 90.1 |
| **CL** | 3.8<sup>a</sup> | 92.6 | 3.8<sup>ab</sup> | 96.3 | 3.5<sup>abc</sup> | 92.6 | 3.6<sup>ab</sup> | 100 | 3.7<sup>ab</sup> | 88.9 | 4.1<sup>a</sup> | 100 | 3.7 | 95.1 |
| **TH** | 1.1<sup>d</sup> | 7.4 | 1.0<sup>c</sup> | 3.7 | 1.0<sup>e</sup> | 0 | 1.0<sup>e</sup> | 0 | 1.1<sup>c</sup> | 7.4 | 1.0<sup>e</sup> | 0 | 1 | 3.1 |
| **FO** | 3.0<sup>c</sup> | 88.9 | 3.0<sup>b</sup> | 77.8 | 2.6<sup>d</sup> | 66.7 | 3.5<sup>abcd</sup> | 92.6 | 3.5<sup>ab</sup> | 92.6 | 3.4<sup>abc</sup> | 92.6 | 3.2 | 85.2 |
| **FV** | 3.3<sup>abc</sup> | 96.3 | 3.3<sup>ab</sup> | 85.2 | 2.9<sup>cd</sup> | 74.1 | 3.3<sup>abcd</sup> | 85.2 | 3.5<sup>ab</sup> | 92.6 | 3.0<sup>cd</sup> | 74.1 | 3.2 | 84.6 |
| **FS** | 2.9<sup>c</sup> | 81.5 | 3.6<sup>ab</sup> | 100 | 3.5<sup>abc</sup> | 96.3 | 2.7<sup>d</sup> | 77.8 | 3.1<sup>b</sup> | 77.8 | 2.7<sup>d</sup> | 77.8 | 3.1 | 85.2 |
| **MP** | 3.7<sup>ab</sup> | 96.3 | 3.9<sup>a</sup> | 96.3 | 3.5<sup>abc</sup> | 92.6 | 3.5<sup>abc</sup> | 100 | 3.8<sup>ab</sup> | 100 | 3.7<sup>abc</sup> | 100 | 3.7 | 97.5 |
| **PES** | 3.9<sup>a</sup> | 100 | 3.6<sup>ab</sup> | 100 | 3.4<sup>abc</sup> | 96.3 | 3.6<sup>ab</sup> | 96.3 | 3.8<sup>ab</sup> | 96.3 | 3.3<sup>bcd</sup> | 100 | 3.6 | 98.1 |
| **PHS** | 3.9<sup>a</sup> | 100 | 4.0<sup>a</sup> | 100 | 3.8<sup>a</sup> | 100 | 3.7<sup>ab</sup> | 100 | 4.1<sup>a</sup> | 100 | 3.2<sup>bcd</sup> | 88.9 | 3.8 | 98.1 |
| **Ckl** | 3.9<sup>a</sup> | 96.3 | 3.7<sup>ab</sup> | 92.6 | 3.6<sup>ab</sup> | 100 | 3b<sup>cd</sup> | 85.2 | 4.2<sup>a</sup> | 100 | 3.9<sup>ab</sup> | 100 | 3.7 | 95.7 |
| **Cont.** | 1.2<sup>d</sup> | 29.6 | 1.2<sup>c</sup> | 14.8 | 1.0<sup>e</sup> | 3.7 | 1.1<sup>e</sup> | 7.4 | 1.2<sup>c</sup> | 18.5 | 1.1<sup>e</sup> | 11.1 | 1.1 | 14.2 |
| **Mean** | 3.2 | 83.3 | 3.2 | 81.5 | 3 | 79.1 | 3 | 79.4 | 3.3 | 81.2 | 3 | 75.9 | 3.1 | 80.1 |

Mean followed by the same letter within a column are not significantly different ( $p < 0.05$ ), according to Duncan's multiple range test.
Pathogen key: AS- *Aspergillus* sp, BS- *Botrytis* sp, BT- *Botryodiplodia theobromae*, CG- *Colletotrichum gloesporioides*, CL- *Cuvularia lunata*, TH- *Trichoderma harzanium*. FO- *Fusarium oxysporum*, FV- *Fusarium verticilloides*, FS- *Fusarium solani*, MP- *Macrophomina phaseolina*, PES-*Pestalotia* sp, PHS- *Phoma* sp, CKL- Cocktail

**Table 5.**
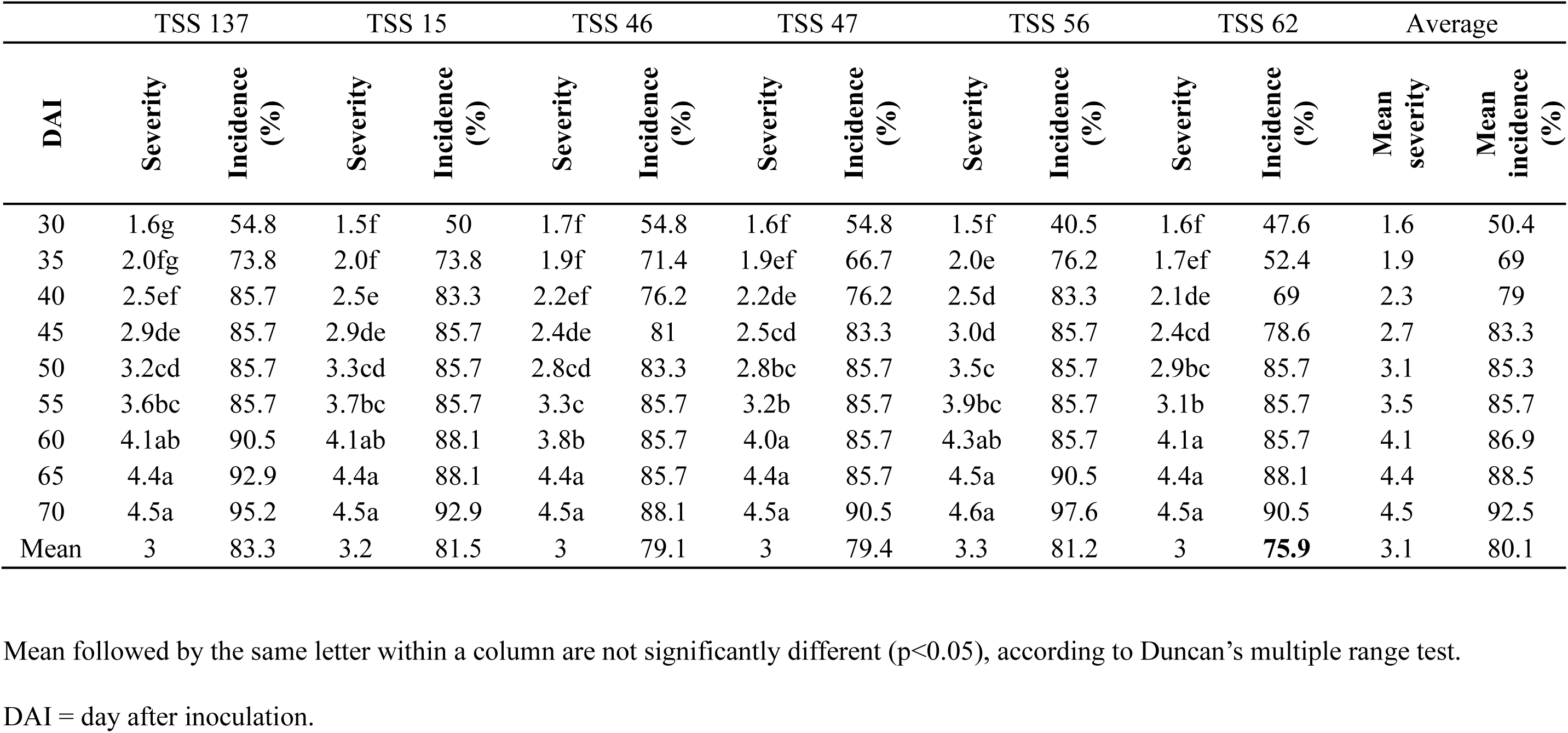
Time course of incidence and disease severity on AYB lines inoculated with fungi under greenhouse conditions.

## 4. DISCUSSION

Fungal diseases represent major constraints to agricultural production, especially in West Africa, where control methods are either not available, not affordable, or not readily accessible by farmers. Effective surveillance of plant diseases usually identifies potential pest problems before they create major crop losses, while permitting sufficient time for mitigation strategies to be developed, tested, and implemented (McCallum et al., 2021).

Field surveys of AYB fungal diseases conducted in SW Nigeria showed incidence of six fungal diseases *viz:* tiny spot, brown spot, brown spot with yellow halo, leaf blight, and brown spot-on pods and with necrotic lesion. These diseases were prevalent and widely distributed in all the surveyed areas in this study regardless of agroecology, altitude, field size, sowing date, cropping system and growth stage. The distribution of these diseases in all surveyed areas might be either due to the environmental conditions which favor development of the disease and/or due to presence of diversified causative pathogens across different AYB growing areas in SW Nigeria. Furthermore, disease incidence may be due to the farming practices adopted by smallholder farmers in the area; for instance the use of poor quality farmer-saved seed lack of crop rotation due to land scarcity, and poor management practices exacerbated by conducive environment conditions for disease development (Njingulula et al., 2014). The higher disease incidence was observed in Oyo and Osun states than other states this could be attributed to the high humidity conditions in these areas. Conversely, the lower occurrence of the disease in Ekiti and Ondo states may reflect the dry and hot conditions in the area when the survey was conducted (2018). This would align with the suggested requirement for high moisture (95-100 % relative humidity) and temperatures (20-25 °C) for the successful infection of the host and sporulation of these pathogens (Liebenberg and Pretorius, 1997). Additionally, there were differences in disease incidence with AYB growth stage. Mean disease incidence at maturity stage for leaf and pod were 56.67 % greater (albeit with greater variation) than early stage at 36.67 %. This would align with a study which have found an association between angular leaf spot disease intensity and common bean growth stages (Kijana et al., 2017). This could also reflect and interaction with developmental changes in the host and the environment which could favor certain infections. Similarly, found higher disease incidences of flower bud infections on 20 accessions of AYB during the wet season (Afolabi et al., 2019). Additionally, a change in agronomic practices might have conducive to the infections at a later AYB developmental stage.

Whatever the conditions that influence disease incidence, any controls mechanisms should focus on identifiable pathogens. In this study, fourteen fungi species belonging to twelve genera were found to be associated with the AYB leaf and pod fungi diseases. Our identified fungal strains (Fig. S1), were similar to those reported by Afolabi *et al*. (2019) in flower bud and pod rot of AYB namely *Fusarium verticilloides, Mucor spp, Fusarium oxysporum, Rhizopus spp, Botryodiplodia spp, Penicillium spp, Aspergillus niger, Macrophomina spp, Curvularia spp, Penicillium oxalicum, Pestalotia spp, Cercospora spp, Pythium spp, Colletotrichum spp and Phomopsis spp.* Although, for logistic reasons, our survey was only conducted over a single year, our field experiments with various genotypes at a single site isolated the same species (Table S1) so further suggested that these were the major fungal pathogens in SW Nigeria. In particular, we found that *Didymella* and *Colletotrichum* were frequently isolated pathogens at early and maturity stages respectively. In this present study, *Phoma/ Didymella* was isolated in all the States of SW Nigeria that were visited. The symptoms of *Didymella* infections range from leaf blight with necrotic spots and chlorotic halos to wilting. Typical blight symptoms include purple-brown irregular flecks on lower leaves, stems, and tendrils, and deep necrotic lesions (Davidson, 2012). Species of *Colletotrichum* are commonly encountered fungal pathogens associated with fruits and a wide range of field crops and produce anthracnose symptoms and leaf spots, rots and seedling infections (Hyde et al., 2009, Cannon et al., 2012). It causes also considerable damage to large number of crops such as legumes, cereals and coffee (Bailey and Jeger, 1992, Lenné, 1992). The symptoms of *Colletotrichum* species appear as small, dark lesions appear on leaves, fruits and flowers of the infected plant which finally produce concentric ring pattern. Taken together, *Phoma/ Didymella* and *Colletotrichum* must be consider major targets in controlling disease in AYB, either through fungicides or developing resistant germplasm. Our field experiments targeted landraces and IITA- donated genotypes which appeared to show reduced disease incidence (Fig. 6) and these could be important in developing field resistance to infection.

It was also relevant that *Fusarium* species were prominent in our AYB-derived samples. *Fusarium* species are regularly associated with decline and yield reduction in the bean growing areas. Characteristic disease symptoms of *Fusarium* species include vascular browning, leaf epinasty, stunting, progressive wilting, defoliation and lastly plant death (Agrios, 2005, Michielse and Rep, 2009). Roots and stems develop a dark-brown discoloration of xylem tissues that can be seen when they are split vertically or cross-sectioned (Jiménez-Díaz et al., 2015) and wilting appears involved gels and gums, blocking or plugging the vessels and leading to symptoms resembling water stress (Pietro et al., 2003). In this study, *Fusarium* spp were isolated from AYB at both early and maturity stages often showing similar symptoms to those just described.

An important part of our work was to confirm that the fungal strains that we had isolated were indeed pathogenic. Therefore, representative strains of most of the tentatively identified genera were tested to fulfil Koch’s postulates under greenhouse conditions. These focused on 12 isolates isolated from 5 states of SW Nigeria and all were pathogenic except *Trichoderma*. The isolates were highly variable in respect of their virulence. The pathogenicity test showed that highest mean percentage incidence (98.1%) and severity (3.8) were caused by *Didymella*. This agrees with previous studies that indicated the pathogenicity of *Didymella americana* on soybean, corn (Aveskamp et al., 2009), wheat (Perelló and Moreno, 2005) and lima bean (Gorny et al., 2016). This result also corroborates with findings of (Singh et al., 2018) who reported the pathogenic of two isolates of *B. theobromae* on Tree bean. The mean severity of *F. oxysporum, is* 3.2 This pathogen has been reported to cause blight or wilt disease in soybean, and has been reported as one of the most destructive diseases of soybean (Hashem et al., 2008, Fayzalla et al., 2009), the pathogen can affect soybeans at any stage of development (Ferrant and Carroll, 1981).

Taken together, our results suggest that any AYB disease control strategies in South Western Africa should focus on *Didymella, Colletotrichum* and *Fusarium.* These strategies should include epidemiological monitoring on field sites (as initiated in this study), improving farm management practices, assessments of fungicide susceptibility and also the development of resistant AYB germplasm possibly including the genotypes that we have targeted.

## Acknowledgements

Thanks go to International Institute of Tropical Agriculture (IITA), Nigeria for providing some of the seed used in this study.

The data that support the findings of this study are available from the corresponding author upon reasonable request.

---

**Figure S1: Identification of Microorganisms of African Yam Bean**

**Fig. S2.** Occurrence (%) of fungi strains at early and mature stages of AYB Pathogen Key : C.C- *Choanephora cucurbitarium*, T.SPP- *Trichoderma* spp, C.L- *Curvularia lunat*a, A.N-A*spergillus* sp*, B.SPP-Botrytis sp, C.G- Collectrotricum gloesporoides, M.P-Macrophomina phaseolina, P.SPP-Pestalotia sp, Phoma* sp, F.V- *Fusarium verticolloides, F.O -F. oxysporum, F.S- F. solani, N.SPP-Nigrospora spp* and B.T- *Botrydipolodia theobromae*

**Fig. S3.** (A) Disease incidence and (B) severity in a field trial of African Yam bean

