## Supplementary material for "Identifying the fungal diseases of African Yam Bean (*Sphenostylis stenocarpa* [Hochst ex. A. Rich.] Harms) and their occurrence in South-West Nigeria": Fig. S1

### Slide 1
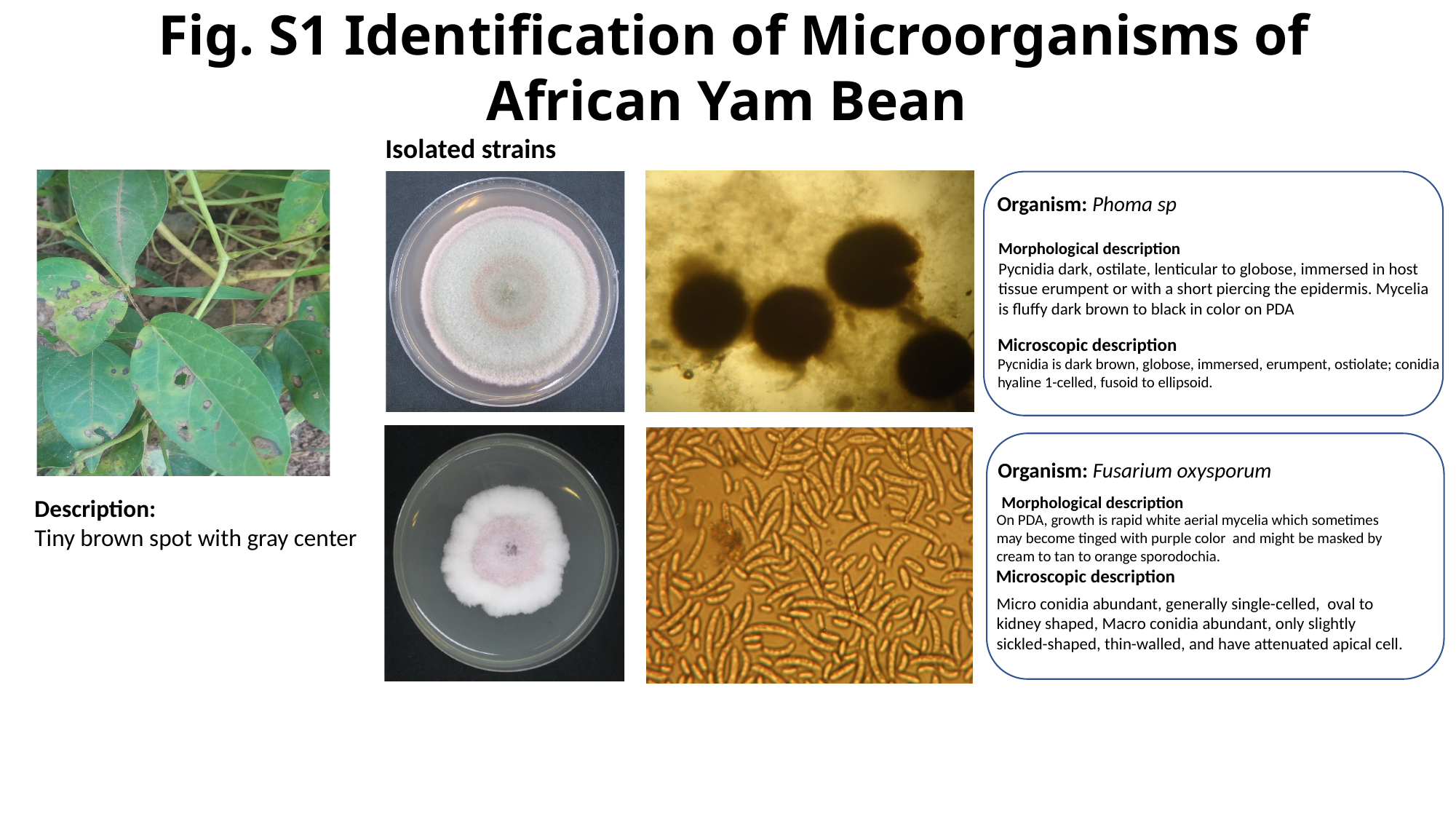

Fig. S1 Identification of Microorganisms of African Yam Bean
Isolated strains
Organism: Phoma sp
Morphological description
Pycnidia dark, ostilate, lenticular to globose, immersed in host tissue erumpent or with a short piercing the epidermis. Mycelia is fluffy dark brown to black in color on PDA
Microscopic description
Pycnidia is dark brown, globose, immersed, erumpent, ostiolate; conidia hyaline 1-celled, fusoid to ellipsoid.
Conidia/spores
Organism: Fusarium oxysporum
Morphological description
Description:
Tiny brown spot with gray center
On PDA, growth is rapid white aerial mycelia which sometimes may become tinged with purple color and might be masked by cream to tan to orange sporodochia.
Microscopic description
Micro conidia abundant, generally single-celled, oval to kidney shaped, Macro conidia abundant, only slightly sickled-shaped, thin-walled, and have attenuated apical cell.

### Slide 2
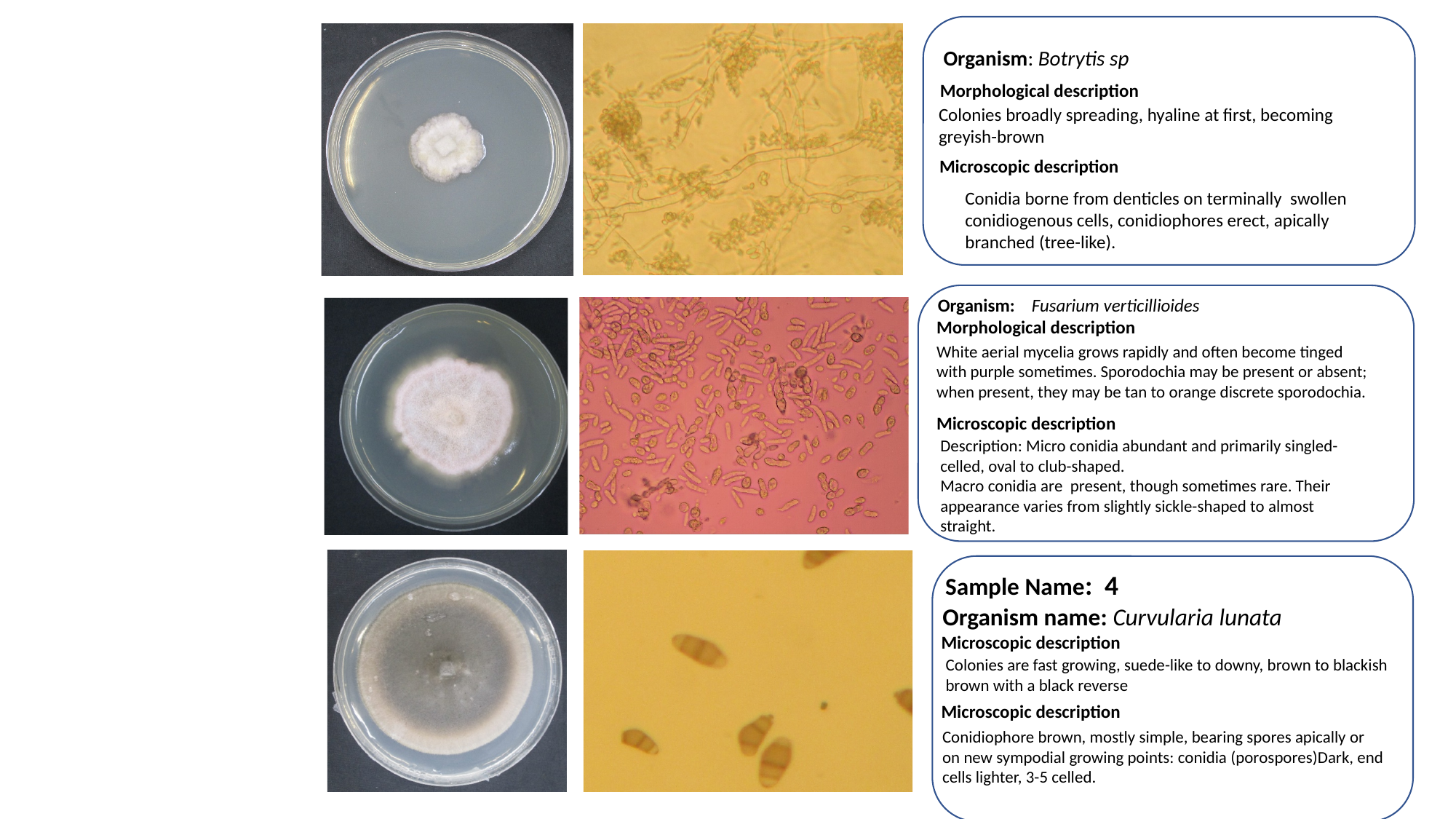

Organism: Botrytis sp
Morphological description
Colonies broadly spreading, hyaline at first, becoming greyish-brown
Microscopic description
Conidia borne from denticles on terminally swollen conidiogenous cells, conidiophores erect, apically branched (tree-like).
Organism: Fusarium verticillioides
Morphological description
White aerial mycelia grows rapidly and often become tinged with purple sometimes. Sporodochia may be present or absent; when present, they may be tan to orange discrete sporodochia.
Microscopic description
Description: Micro conidia abundant and primarily singled-celled, oval to club-shaped.
Macro conidia are present, though sometimes rare. Their appearance varies from slightly sickle-shaped to almost straight.
Sample Name: 4
Organism name: Curvularia lunata
Microscopic description
Colonies are fast growing, suede-like to downy, brown to blackish
brown with a black reverse
Microscopic description
Conidiophore brown, mostly simple, bearing spores apically or on new sympodial growing points: conidia (porospores)Dark, end cells lighter, 3-5 celled.

### Slide 3
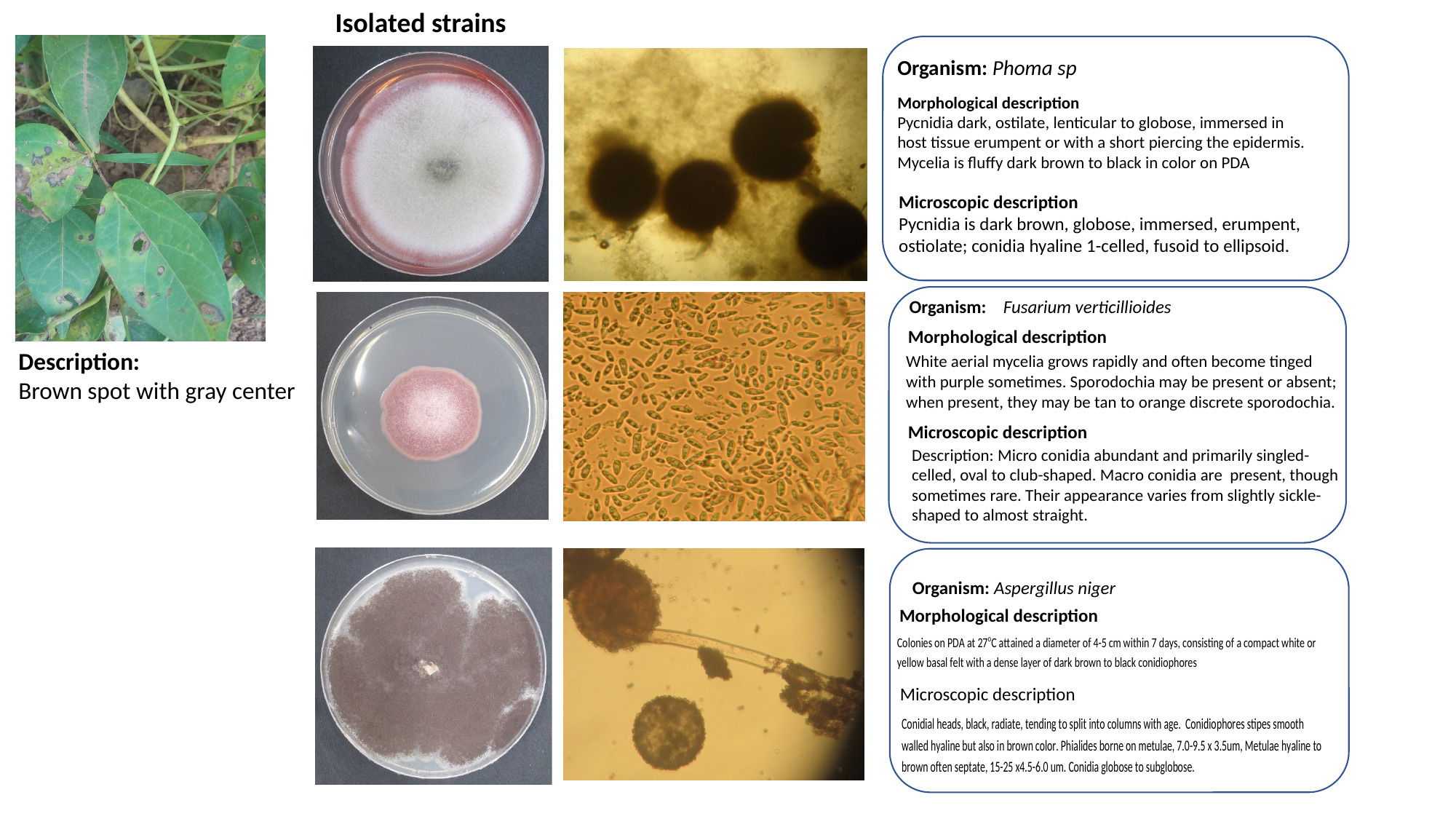

Isolated strains
Organism: Phoma sp
Morphological description
Pycnidia dark, ostilate, lenticular to globose, immersed in
host tissue erumpent or with a short piercing the epidermis.
Mycelia is fluffy dark brown to black in color on PDA
Microscopic description
Pycnidia is dark brown, globose, immersed, erumpent, ostiolate; conidia hyaline 1-celled, fusoid to ellipsoid.
Organism: Fusarium verticillioides
Morphological description
Description:
Brown spot with gray center
White aerial mycelia grows rapidly and often become tinged with purple sometimes. Sporodochia may be present or absent; when present, they may be tan to orange discrete sporodochia.
Microscopic description
Description: Micro conidia abundant and primarily singled-celled, oval to club-shaped. Macro conidia are present, though sometimes rare. Their appearance varies from slightly sickle-shaped to almost straight.
 Organism: Aspergillus niger
Morphological description
Microscopic description

### Slide 4
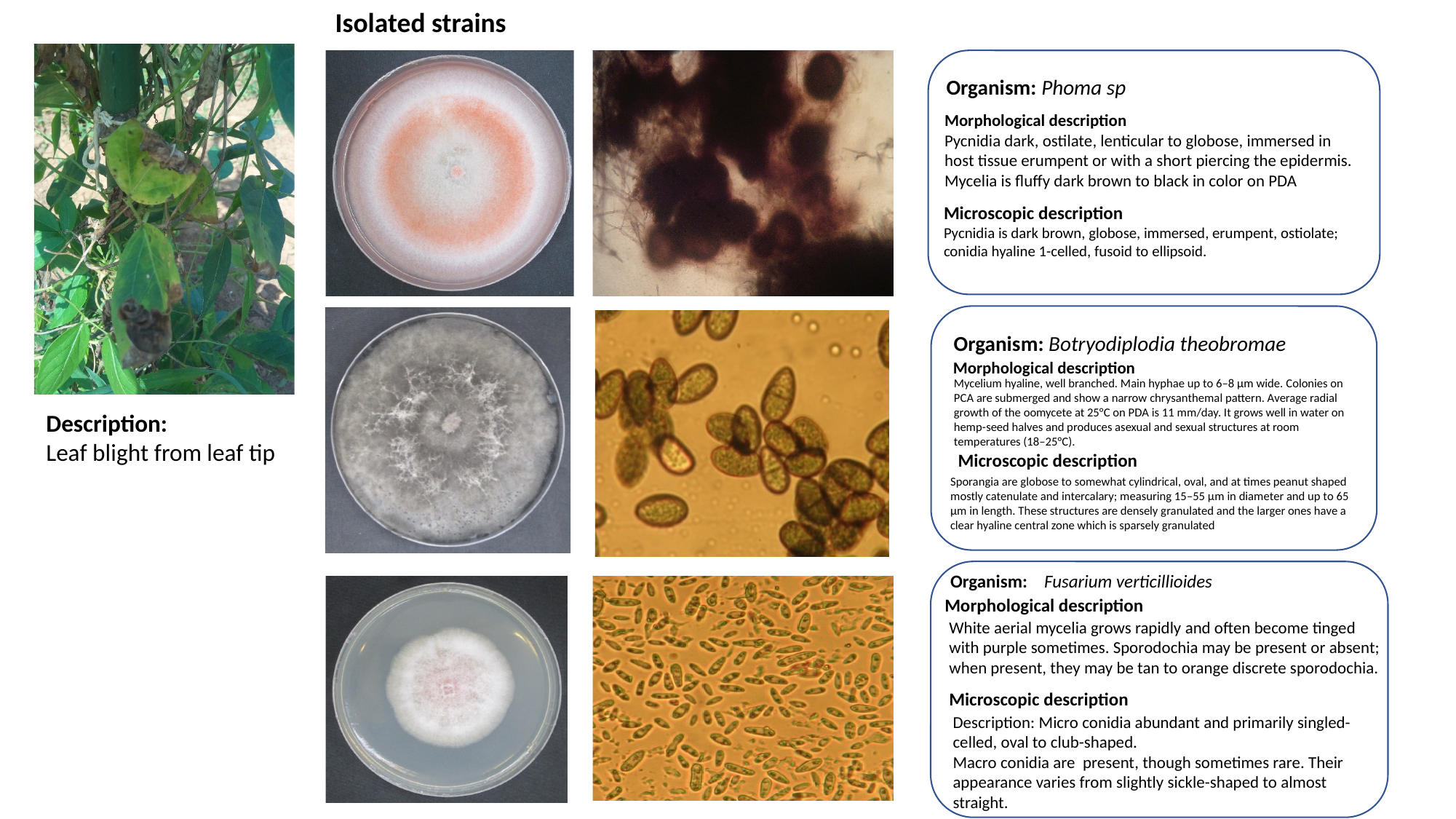

Isolated strains
Organism: Phoma sp
Morphological description
Pycnidia dark, ostilate, lenticular to globose, immersed in host tissue erumpent or with a short piercing the epidermis. Mycelia is fluffy dark brown to black in color on PDA
Microscopic description
Pycnidia is dark brown, globose, immersed, erumpent, ostiolate; conidia hyaline 1-celled, fusoid to ellipsoid.
Organism: Botryodiplodia theobromae
Morphological description
Mycelium hyaline, well branched. Main hyphae up to 6–8 μm wide. Colonies on PCA are submerged and show a narrow chrysanthemal pattern. Average radial growth of the oomycete at 25°C on PDA is 11 mm/day. It grows well in water on hemp-seed halves and produces asexual and sexual structures at room temperatures (18–25°C).
Description:
Leaf blight from leaf tip
Microscopic description
Sporangia are globose to somewhat cylindrical, oval, and at times peanut shaped mostly catenulate and intercalary; measuring 15–55 μm in diameter and up to 65 μm in length. These structures are densely granulated and the larger ones have a clear hyaline central zone which is sparsely granulated
Organism: Fusarium verticillioides
Morphological description
White aerial mycelia grows rapidly and often become tinged with purple sometimes. Sporodochia may be present or absent; when present, they may be tan to orange discrete sporodochia.
Microscopic description
Description: Micro conidia abundant and primarily singled-celled, oval to club-shaped.
Macro conidia are present, though sometimes rare. Their appearance varies from slightly sickle-shaped to almost straight.

### Slide 5
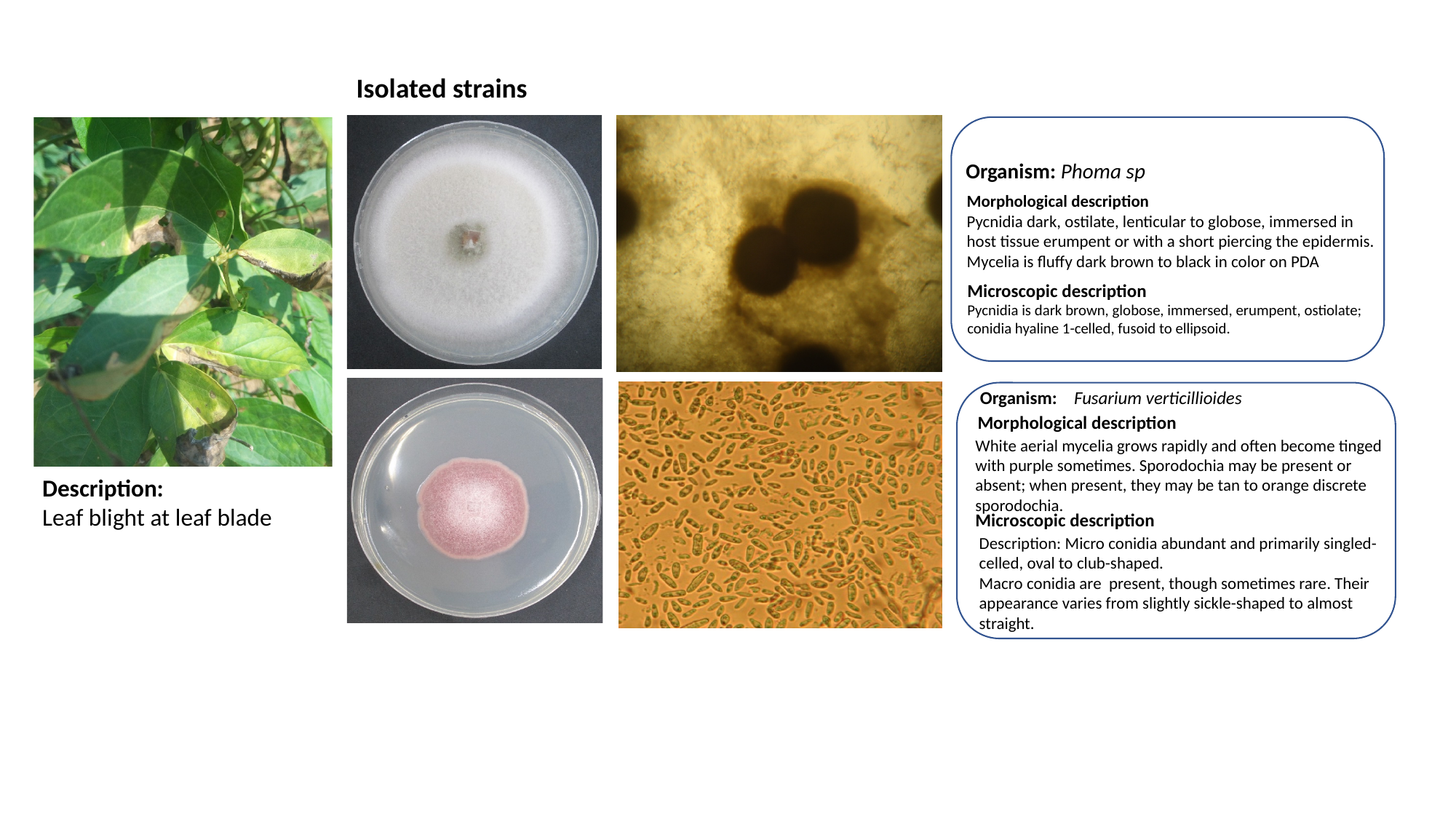

Isolated strains
Organism: Phoma sp
Morphological description
Pycnidia dark, ostilate, lenticular to globose, immersed in
host tissue erumpent or with a short piercing the epidermis.
Mycelia is fluffy dark brown to black in color on PDA
Microscopic description
Pycnidia is dark brown, globose, immersed, erumpent, ostiolate; conidia hyaline 1-celled, fusoid to ellipsoid.
Organism: Fusarium verticillioides
Morphological description
White aerial mycelia grows rapidly and often become tinged with purple sometimes. Sporodochia may be present or absent; when present, they may be tan to orange discrete sporodochia.
Description:
Leaf blight at leaf blade
Microscopic description
Description: Micro conidia abundant and primarily singled-celled, oval to club-shaped.
Macro conidia are present, though sometimes rare. Their appearance varies from slightly sickle-shaped to almost straight.

### Slide 6
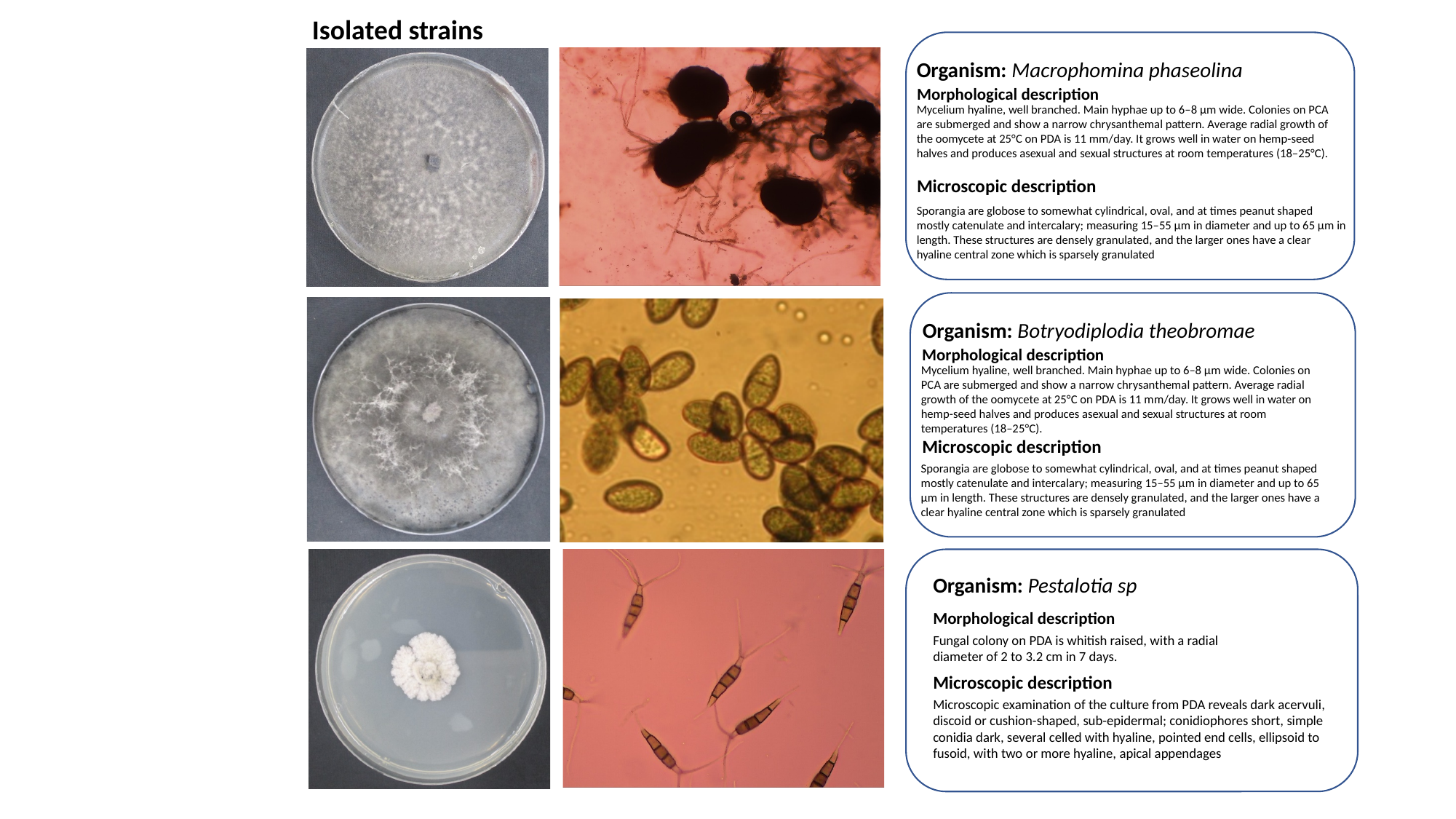

Isolated strains
Organism: Macrophomina phaseolina
Morphological description
Mycelium hyaline, well branched. Main hyphae up to 6–8 μm wide. Colonies on PCA are submerged and show a narrow chrysanthemal pattern. Average radial growth of the oomycete at 25°C on PDA is 11 mm/day. It grows well in water on hemp-seed halves and produces asexual and sexual structures at room temperatures (18–25°C).
Microscopic description
Sporangia are globose to somewhat cylindrical, oval, and at times peanut shaped mostly catenulate and intercalary; measuring 15–55 μm in diameter and up to 65 μm in length. These structures are densely granulated, and the larger ones have a clear hyaline central zone which is sparsely granulated
Organism: Botryodiplodia theobromae
Morphological description
Mycelium hyaline, well branched. Main hyphae up to 6–8 μm wide. Colonies on PCA are submerged and show a narrow chrysanthemal pattern. Average radial growth of the oomycete at 25°C on PDA is 11 mm/day. It grows well in water on hemp-seed halves and produces asexual and sexual structures at room temperatures (18–25°C).
Microscopic description
Sporangia are globose to somewhat cylindrical, oval, and at times peanut shaped mostly catenulate and intercalary; measuring 15–55 μm in diameter and up to 65 μm in length. These structures are densely granulated, and the larger ones have a clear hyaline central zone which is sparsely granulated
Organism: Pestalotia sp
Morphological description
Fungal colony on PDA is whitish raised, with a radial
diameter of 2 to 3.2 cm in 7 days.
Microscopic description
Microscopic examination of the culture from PDA reveals dark acervuli, discoid or cushion-shaped, sub-epidermal; conidiophores short, simple conidia dark, several celled with hyaline, pointed end cells, ellipsoid to fusoid, with two or more hyaline, apical appendages

### Slide 7
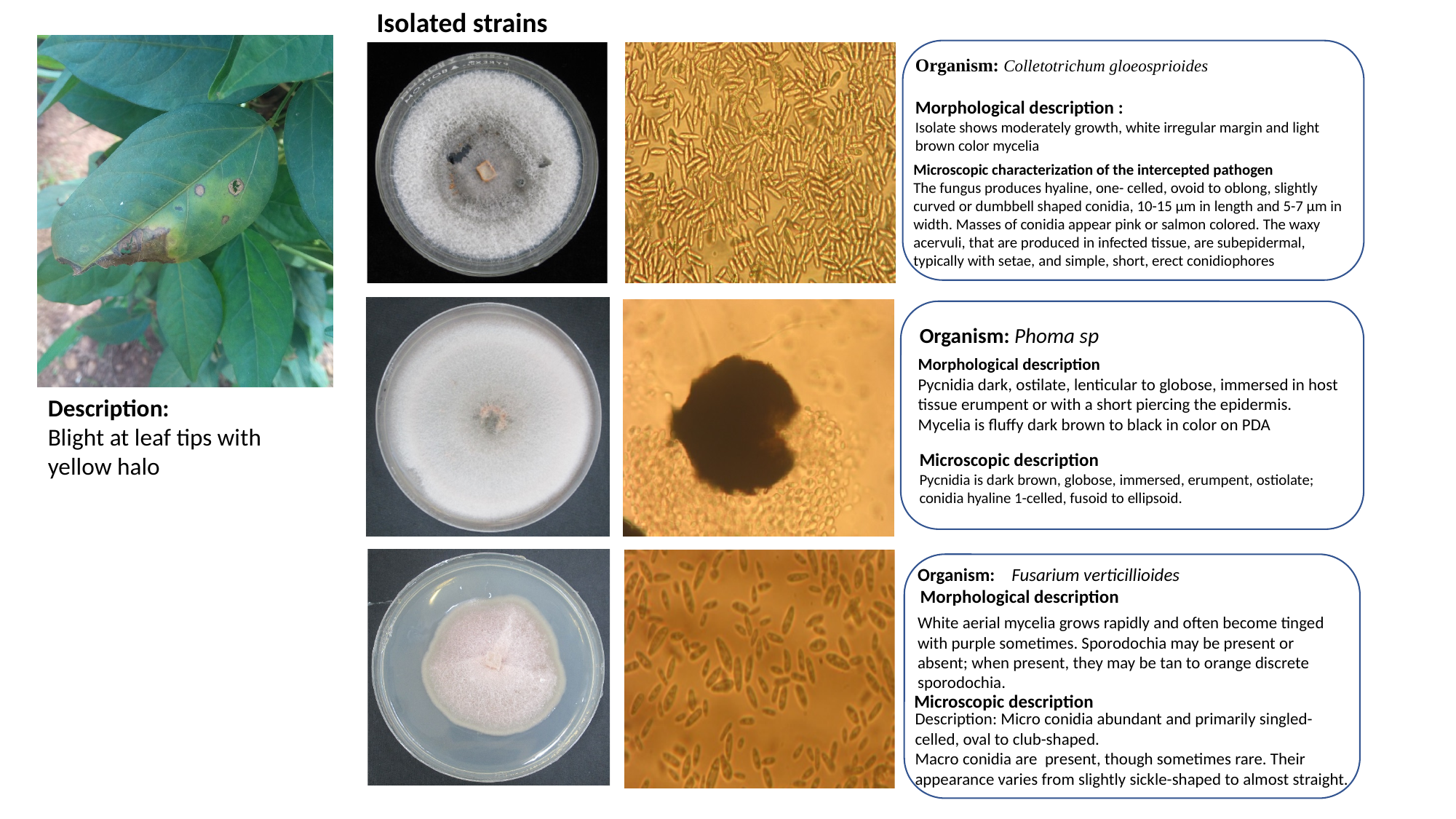

Isolated strains
Organism: Colletotrichum gloeosprioides
Morphological description :
Isolate shows moderately growth, white irregular margin and light brown color mycelia
Microscopic characterization of the intercepted pathogen
The fungus produces hyaline, one- celled, ovoid to oblong, slightly curved or dumbbell shaped conidia, 10-15 µm in length and 5-7 µm in width. Masses of conidia appear pink or salmon colored. The waxy acervuli, that are produced in infected tissue, are subepidermal, typically with setae, and simple, short, erect conidiophores
Conidia/spores
Organism: Phoma sp
Morphological description
Pycnidia dark, ostilate, lenticular to globose, immersed in host tissue erumpent or with a short piercing the epidermis. Mycelia is fluffy dark brown to black in color on PDA
Description:
Blight at leaf tips with
yellow halo
Microscopic description
Pycnidia is dark brown, globose, immersed, erumpent, ostiolate; conidia hyaline 1-celled, fusoid to ellipsoid.
Organism: Fusarium verticillioides
Morphological description
White aerial mycelia grows rapidly and often become tinged with purple sometimes. Sporodochia may be present or absent; when present, they may be tan to orange discrete sporodochia.
Microscopic description
Description: Micro conidia abundant and primarily singled-celled, oval to club-shaped.
Macro conidia are present, though sometimes rare. Their appearance varies from slightly sickle-shaped to almost straight.

### Slide 8
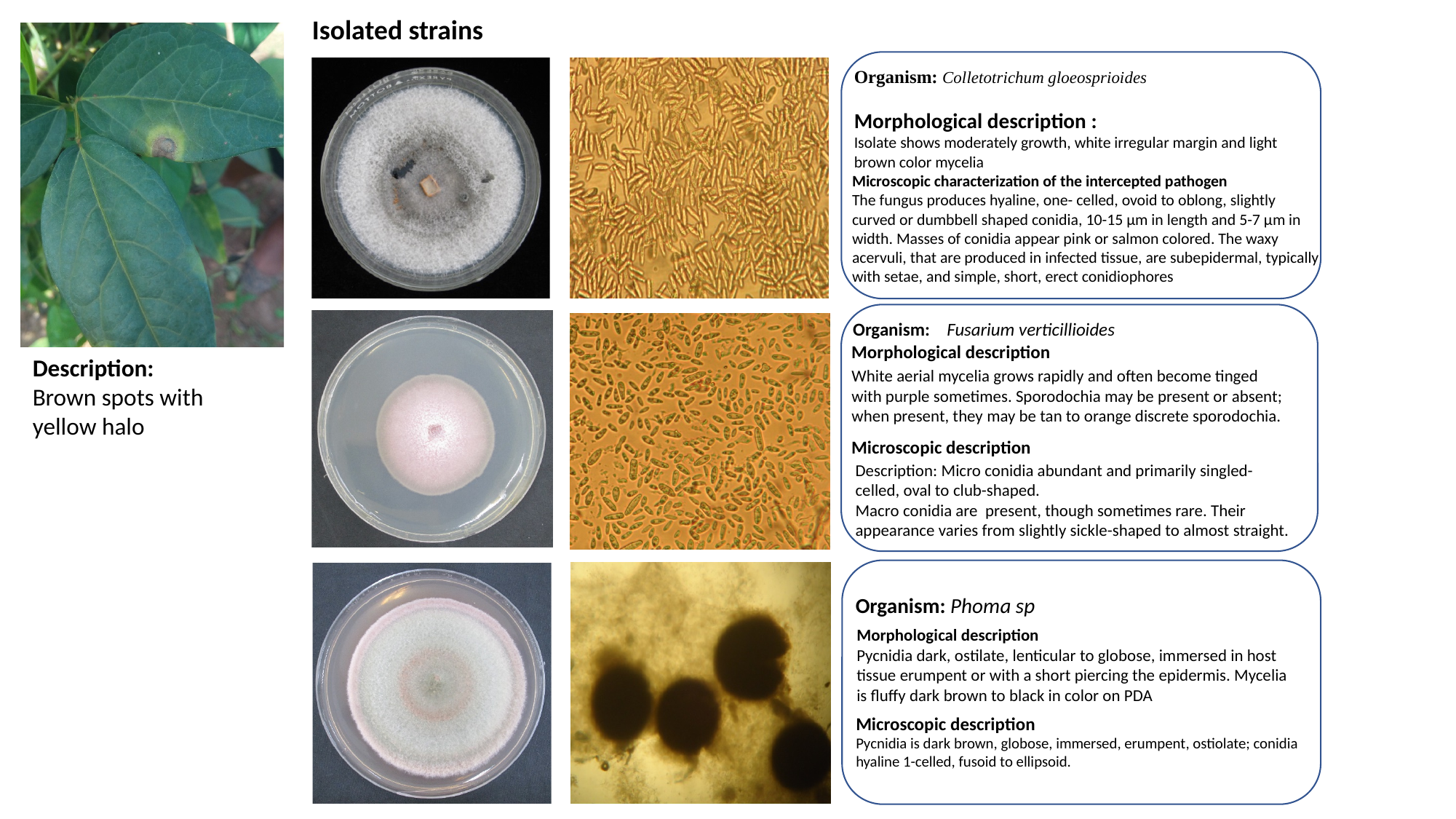

Isolated strains
Organism: Colletotrichum gloeosprioides
Morphological description :
Isolate shows moderately growth, white irregular margin and light brown color mycelia
Microscopic characterization of the intercepted pathogen
The fungus produces hyaline, one- celled, ovoid to oblong, slightly curved or dumbbell shaped conidia, 10-15 µm in length and 5-7 µm in width. Masses of conidia appear pink or salmon colored. The waxy acervuli, that are produced in infected tissue, are subepidermal, typically with setae, and simple, short, erect conidiophores
Conidia/spores
Organism: Fusarium verticillioides
Morphological description
Description:
Brown spots with
yellow halo
White aerial mycelia grows rapidly and often become tinged with purple sometimes. Sporodochia may be present or absent; when present, they may be tan to orange discrete sporodochia.
Microscopic description
Description: Micro conidia abundant and primarily singled-celled, oval to club-shaped.
Macro conidia are present, though sometimes rare. Their appearance varies from slightly sickle-shaped to almost straight.
Organism: Phoma sp
Morphological description
Pycnidia dark, ostilate, lenticular to globose, immersed in host tissue erumpent or with a short piercing the epidermis. Mycelia is fluffy dark brown to black in color on PDA
Microscopic description
Pycnidia is dark brown, globose, immersed, erumpent, ostiolate; conidia hyaline 1-celled, fusoid to ellipsoid.

### Slide 9
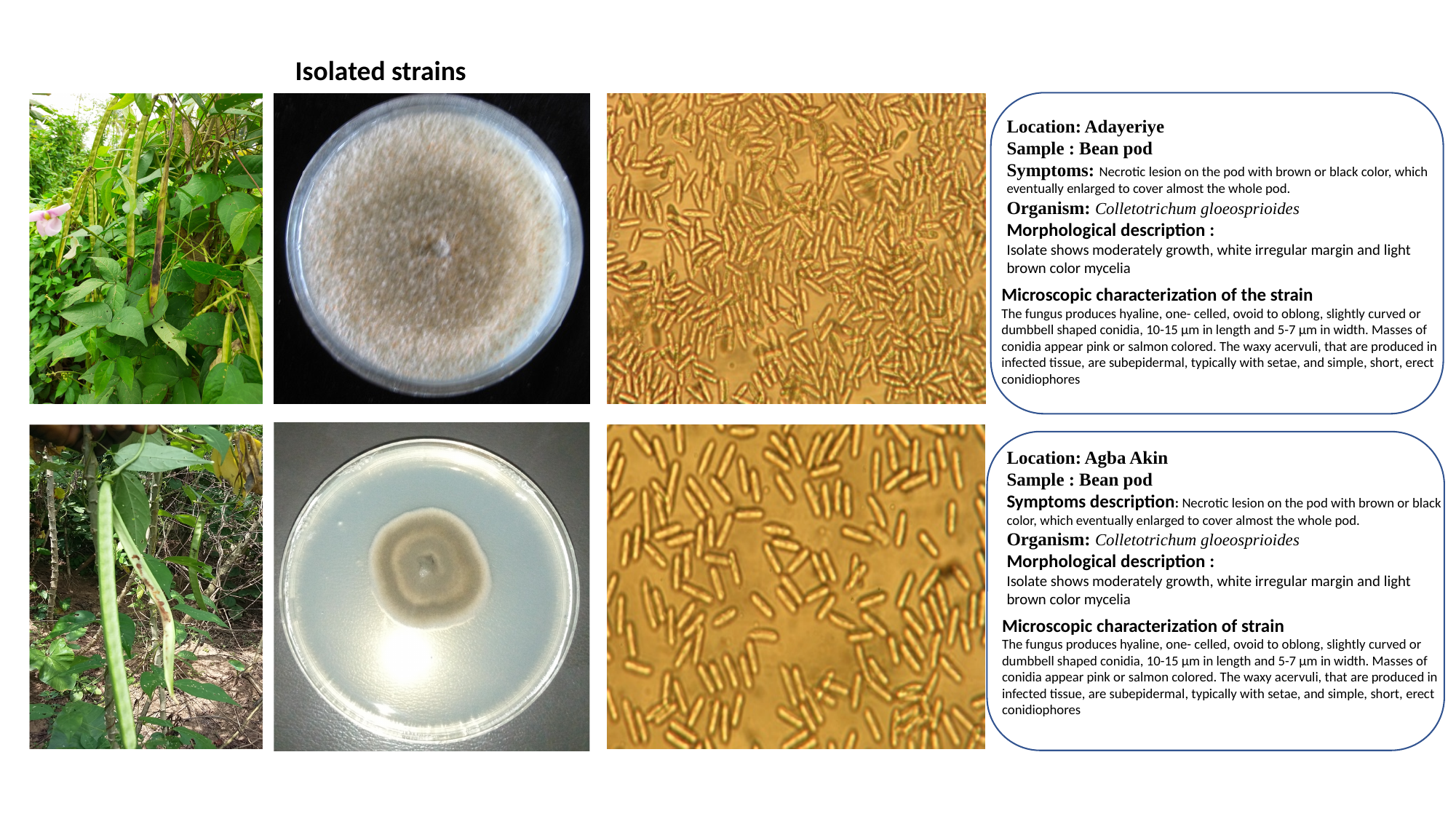

Isolated strains
Location: Adayeriye
Sample : Bean pod
Symptoms: Necrotic lesion on the pod with brown or black color, which eventually enlarged to cover almost the whole pod.
Organism: Colletotrichum gloeosprioides
Morphological description :
Isolate shows moderately growth, white irregular margin and light brown color mycelia
Microscopic characterization of the strain
The fungus produces hyaline, one- celled, ovoid to oblong, slightly curved or dumbbell shaped conidia, 10-15 µm in length and 5-7 µm in width. Masses of conidia appear pink or salmon colored. The waxy acervuli, that are produced in infected tissue, are subepidermal, typically with setae, and simple, short, erect conidiophores
Location: Agba Akin
Sample : Bean pod
Symptoms description: Necrotic lesion on the pod with brown or black color, which eventually enlarged to cover almost the whole pod.
Organism: Colletotrichum gloeosprioides
Morphological description :
Isolate shows moderately growth, white irregular margin and light brown color mycelia
Microscopic characterization of strain
The fungus produces hyaline, one- celled, ovoid to oblong, slightly curved or dumbbell shaped conidia, 10-15 µm in length and 5-7 µm in width. Masses of conidia appear pink or salmon colored. The waxy acervuli, that are produced in infected tissue, are subepidermal, typically with setae, and simple, short, erect conidiophores

### Slide 10
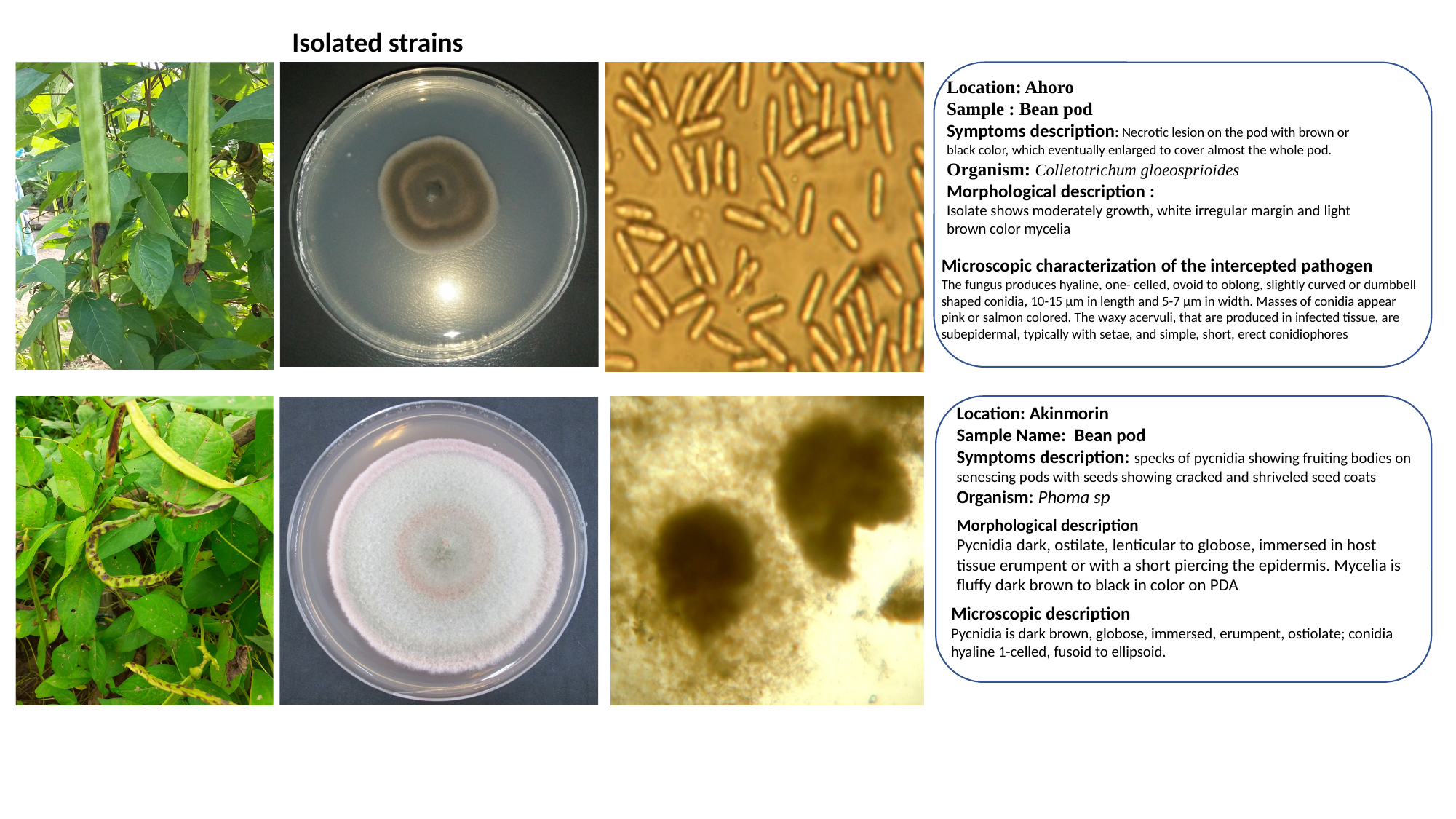

Isolated strains
Location: Ahoro
Sample : Bean pod
Symptoms description: Necrotic lesion on the pod with brown or black color, which eventually enlarged to cover almost the whole pod.
Organism: Colletotrichum gloeosprioides
Morphological description :
Isolate shows moderately growth, white irregular margin and light brown color mycelia
Microscopic characterization of the intercepted pathogen
The fungus produces hyaline, one- celled, ovoid to oblong, slightly curved or dumbbell shaped conidia, 10-15 µm in length and 5-7 µm in width. Masses of conidia appear pink or salmon colored. The waxy acervuli, that are produced in infected tissue, are subepidermal, typically with setae, and simple, short, erect conidiophores
Location: Akinmorin
Sample Name: Bean pod
Symptoms description: specks of pycnidia showing fruiting bodies on senescing pods with seeds showing cracked and shriveled seed coats
Organism: Phoma sp
Morphological description
Pycnidia dark, ostilate, lenticular to globose, immersed in host tissue erumpent or with a short piercing the epidermis. Mycelia is fluffy dark brown to black in color on PDA
Microscopic description
Pycnidia is dark brown, globose, immersed, erumpent, ostiolate; conidia hyaline 1-celled, fusoid to ellipsoid.

### Slide 11
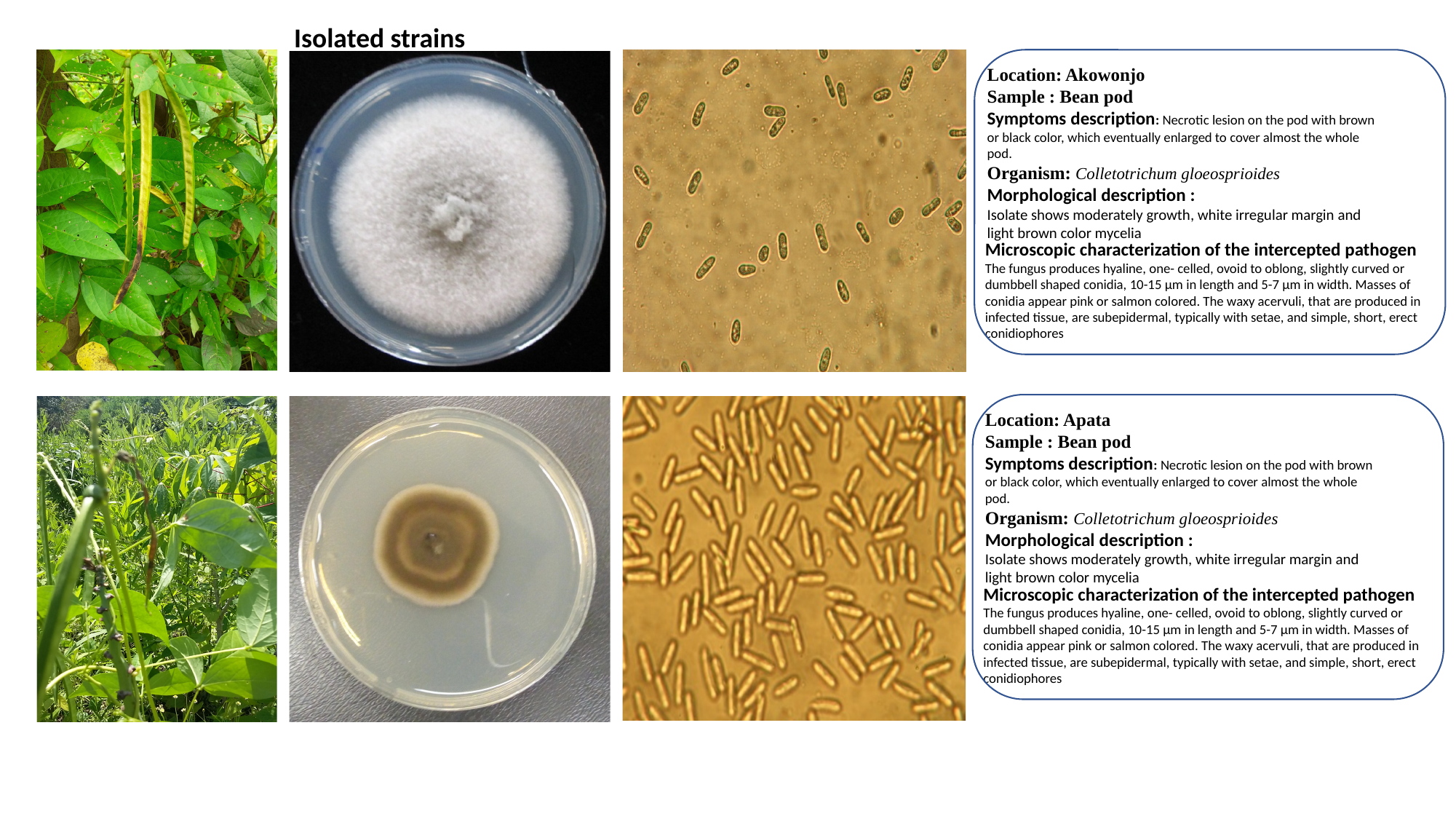

Isolated strains
Location: Akowonjo
Sample : Bean pod
Symptoms description: Necrotic lesion on the pod with brown or black color, which eventually enlarged to cover almost the whole pod.
Organism: Colletotrichum gloeosprioides
Morphological description :
Isolate shows moderately growth, white irregular margin and light brown color mycelia
Microscopic characterization of the intercepted pathogen
The fungus produces hyaline, one- celled, ovoid to oblong, slightly curved or dumbbell shaped conidia, 10-15 µm in length and 5-7 µm in width. Masses of conidia appear pink or salmon colored. The waxy acervuli, that are produced in infected tissue, are subepidermal, typically with setae, and simple, short, erect conidiophores
Location: Apata
Sample : Bean pod
Symptoms description: Necrotic lesion on the pod with brown or black color, which eventually enlarged to cover almost the whole pod.
Organism: Colletotrichum gloeosprioides
Morphological description :
Isolate shows moderately growth, white irregular margin and light brown color mycelia
Microscopic characterization of the intercepted pathogen
The fungus produces hyaline, one- celled, ovoid to oblong, slightly curved or dumbbell shaped conidia, 10-15 µm in length and 5-7 µm in width. Masses of conidia appear pink or salmon colored. The waxy acervuli, that are produced in infected tissue, are subepidermal, typically with setae, and simple, short, erect conidiophores

### Slide 12
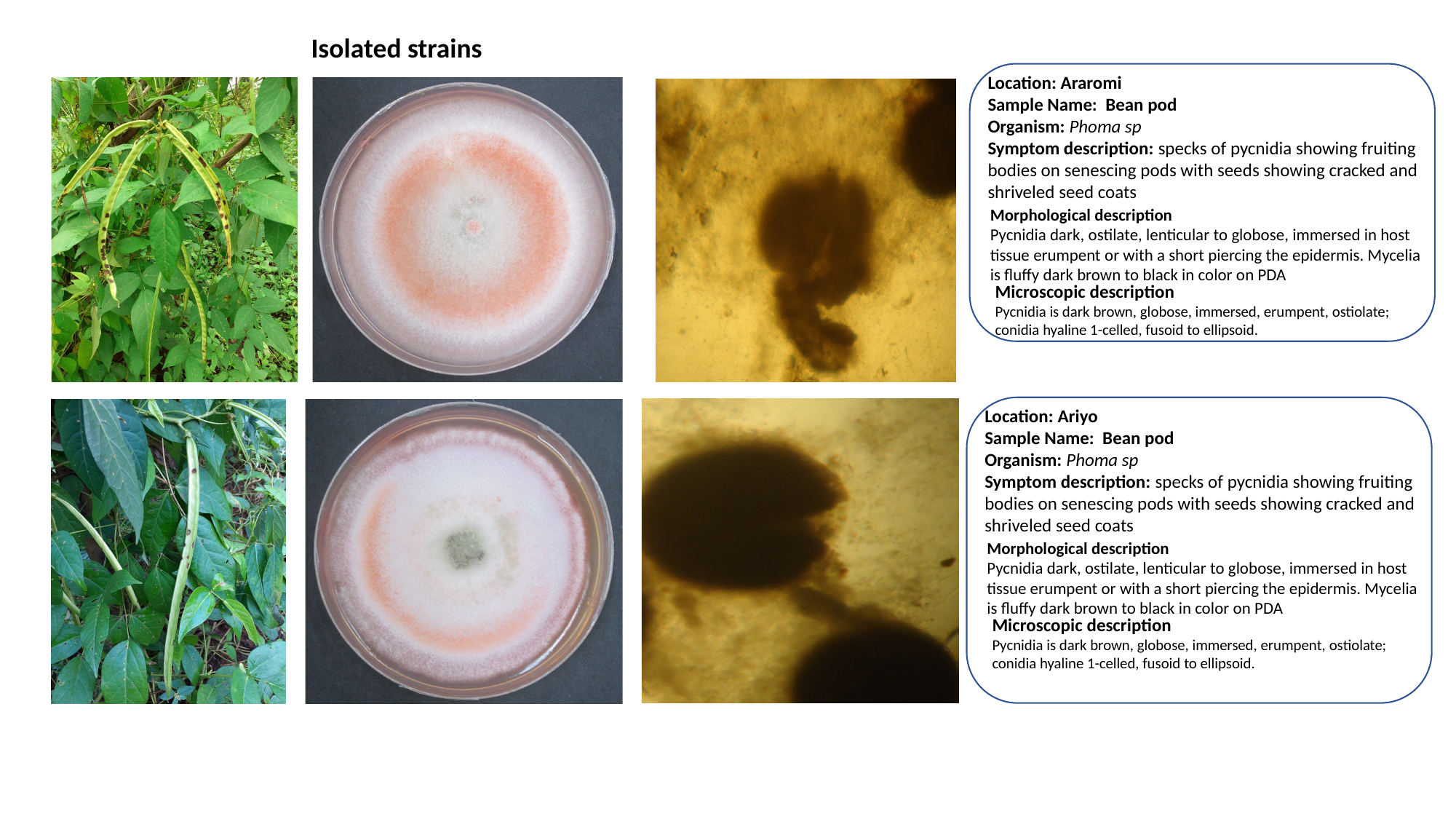

Isolated strains
Location: Araromi
Sample Name: Bean pod
Organism: Phoma sp
Symptom description: specks of pycnidia showing fruiting bodies on senescing pods with seeds showing cracked and shriveled seed coats
Morphological description
Pycnidia dark, ostilate, lenticular to globose, immersed in host tissue erumpent or with a short piercing the epidermis. Mycelia is fluffy dark brown to black in color on PDA
Microscopic description
Pycnidia is dark brown, globose, immersed, erumpent, ostiolate; conidia hyaline 1-celled, fusoid to ellipsoid.
Location: Ariyo
Sample Name: Bean pod
Organism: Phoma sp
Symptom description: specks of pycnidia showing fruiting bodies on senescing pods with seeds showing cracked and shriveled seed coats
Morphological description
Pycnidia dark, ostilate, lenticular to globose, immersed in host tissue erumpent or with a short piercing the epidermis. Mycelia is fluffy dark brown to black in color on PDA
Microscopic description
Pycnidia is dark brown, globose, immersed, erumpent, ostiolate; conidia hyaline 1-celled, fusoid to ellipsoid.
