## Supplementary figures and images for "Identifying the fungal diseases of African Yam Bean (*Sphenostylis stenocarpa* [Hochst ex. A. Rich.] Harms) and their occurrence in South-West Nigeria"

### Fig. S2

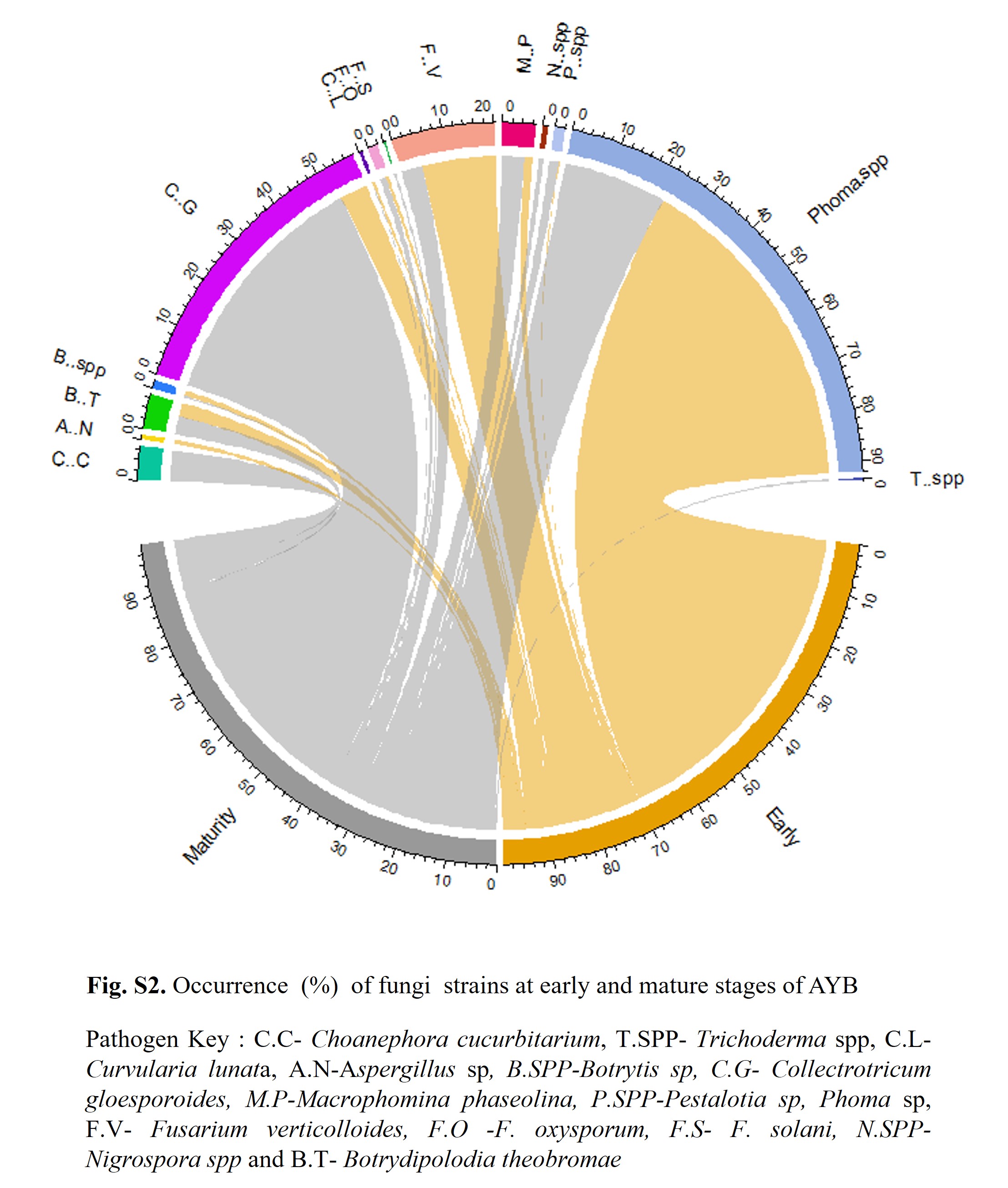

### Fig. S3

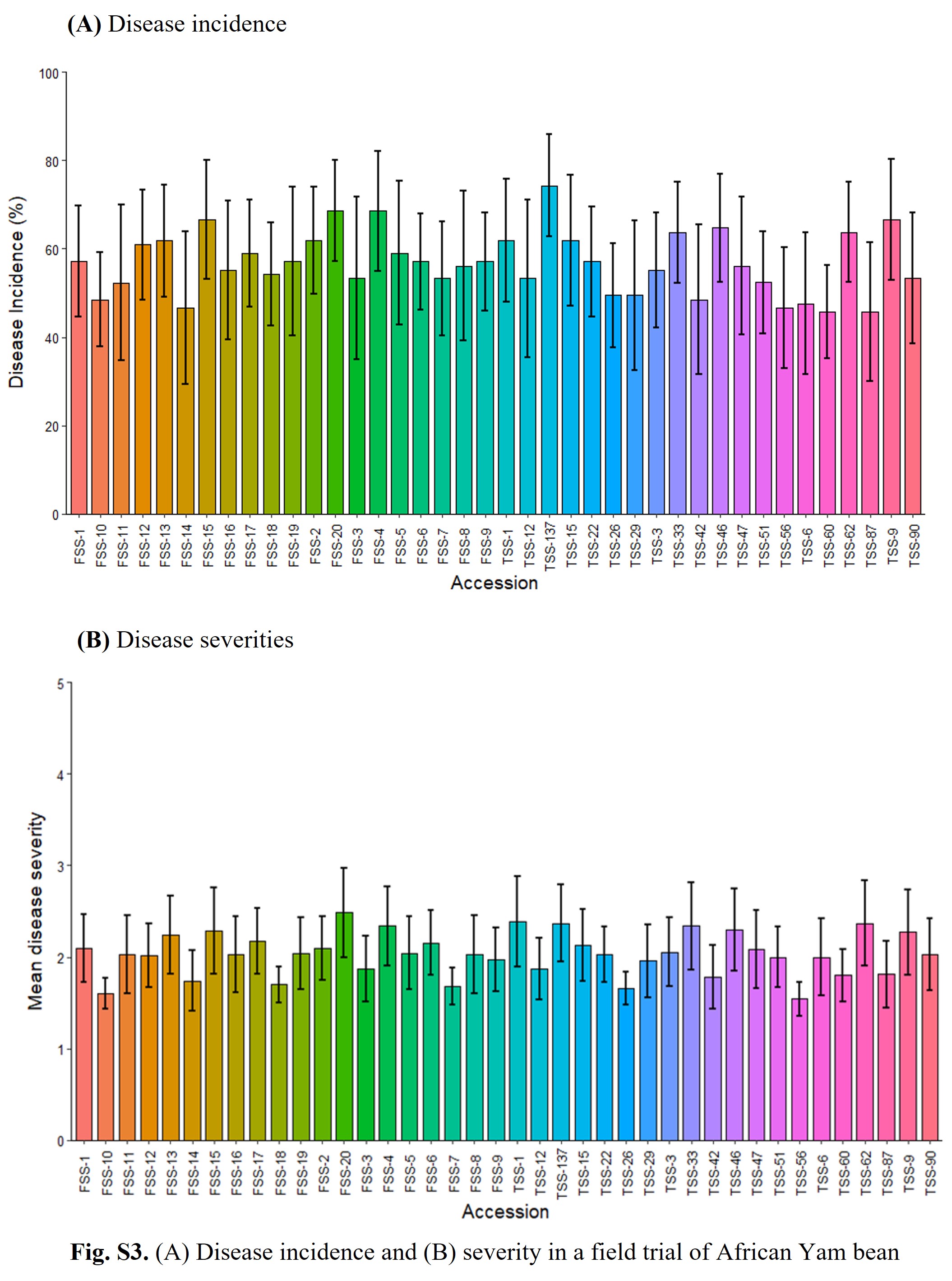
